# Macrophage-derived cholesterol contributes to therapeutic resistance in prostate cancer

**DOI:** 10.1101/2021.03.24.436480

**Authors:** Asmaa El-Kenawi, William Dominguez-Viqueira, Min Liu, Shivanshu Awasthi, Aysenur Keske, KayLee K. Steiner, Leenil Noel, Jasreman Dhillon, Robert J. Gillies, Kosj Yamoah, Xiaoqing Yu, John Koomen, Robert A. Gatenby, Brian Ruffell

**Author notes:** Correspondence, (A.E.), (B.R.). **Contact:** Brian Ruffell, Ph.D., H. Lee Moffitt Cancer Center, 12902 Magnolia Drive SRB-2, Tampa, FL, 33612, Voice.

## Abstract

Tumor-associated macrophages are key immune cells associated with cancer progression. Here we sought to determine the role of macrophages in castration-resistant prostate cancer (CRPC) using a syngeneic model that reflected the mutational landscape of the disease. A transcriptomic analysis of CRPC tumors following macrophage depletion revealed lower molecular signatures for steroid and bile acid synthesis, indicating potential perturbation of cholesterol metabolism. Since cholesterol is the precursor of the five major classes of steroid hormones, we reasoned that macrophages were regulating androgen biosynthesis within the prostate tumor microenvironment. Indeed, macrophage depletion reduced the levels of androgens within prostate tumors and restricted androgen receptor (AR) nuclear localization *in vitro* and *in vivo*. Macrophages were cholesterol rich and had the ability to transfer cholesterol to tumor cells *in vitro*, and AR nuclear translocation was inhibited by activation of Liver X Receptor (LXR)-β, the master regulator of cholesterol homeostasis. Finally, combining macrophage depletion with androgen deprivation therapy increased survival, supporting the therapeutic potential of targeting macrophages in CRPC.

## Introduction

Prostate cancer is the second leading of cancer death in men in the United States. Androgen hormones drive prostate cancer cell proliferation and progression by activating the androgen receptor (AR). Given their tumor-promoting role, reducing androgen levels or signaling by using Androgen Deprivation Therapy (ADT) is the standard approach to treat advanced prostate cancer (1,2). In most cases ADT induces tumor regression; however, resistance eventually develops and the disease relapses as Castration-Resistant Prostate Cancer (CRPC) (3,4). Androgen-AR signaling persists in many cases of CRPC via alterations in the AR gene, deregulation of AR binding-proteins (2) and/or intratumoral synthesis of androgens (5); thus, both androgen synthesis inhibitors and AR inhibitors can extend survival (6). However, while much is known regarding the intrinsic mechanisms of AR reactivation within tumor cells, little is understood regarding the contributions of the tumor microenvironment in therapeutic resistance in CRPC.

Macrophages are the most abundant immune subpopulation in many solid tumors, wherein they adopt a maladaptive phenotype and are commonly known as tumor-associated macrophages (TAMs) (7). In addition to their role in creating the immunosuppressive microenvironment in tumors (8), TAMs can promote tumor growth and resistance to therapy through the release of cytokines and angiogenic mediators (9,10). More recently, TAM-secreted metabolites have emerged as a driver of drug resistance. TAM-derived pyrimidines diminished the efficacy of gemcitabine in pancreatic cancer through molecular competition during drug uptake, while in a model of colon cancer, TAMs were involved in the production of glucocorticoids and impaired the efficacy of immune checkpoint blockade (11). Whether macrophage-tumor cell metabolic interactions dictate therapeutic responses in prostate cancer is unknown.

Here we sought to determine the role of macrophages in CRPC using a murine model that recapitulated the high AR signaling and mutational landscape of prostate cancer. We developed orthotopic systems derived from an inducible, autochthonous model of *Trp53/Pten-deficiency* with or without the most common *ETS* rearrangement, *TMPRSS2/ERG* fusion. We then performed transcriptomic and metabolomic characterization of prostate tumors following macrophage depletion. Combined with efficacy studies and *in vitro* experiments, our data demonstrate that macrophages promote AR signaling and castration resistance via enhanced tumor cell cholesterol influx and androgen biosynthesis.

## Materials and Methods

### Animal models

All mice were maintained in accordance with Institutional Animal Care and Use Committee (IACUC) standards followed by the Moffitt Cancer Research Center (Tampa, Florida). The conditional model of prostate cancer was established by crossing Trp53^loxP^ (Trp53^tm1Brn^/J; JAX 008462), Pten^loxP^ (Pten^tm1Hwu^/J; JAX 006440) and C57BL/6-Tg(Pbsn-cre/Esr1*)14Abch/J mice (JAX 020287), with or without expression of the *TMPRSS2/ERG* fusion gene under control of the Rosa26 promoter and a loxP-flanked stop sequence (B6.129-Gt(ROSA)26Sor^tm1(TMPRSS2/ERG)Key^/J; JAX 024512). The Pbsn-cre/Esr1* transgene allows inducible Cre-mediated recombination in the mouse prostate following ad libitum dosing of 500 mg/kg tamoxifen-containing chow for 4 weeks(12). Dual *Abca1*/*Abcg1*-floxed mice (B6.Cg-*Abca1*^tm1Jp^ *Abcg1*^tm1Tall^/J; JAX 021067) were crossed with (B6.129P2-Lyz2^tm1(cre)Ifo^/J; JAX 004781). Male C57BL/6J (JAX 000664; 8-12 weeks of age) mice purchased from The Jackson Laboratory were utilized for orthotopic prostate cancer models and bone marrow isolation. Mice were treated with 0.6 mg Lupron Depot (AbbVie Inc. North Chicago) by subcutaneous injection. αCSF1 (clone 5A1, BioXCell) was administrated intraperitoneally at 1 mg/mouse, followed by 0.5 mg/mouse every 5 days (13).

### Magnetic resonance Imaging (MRI)

All MRI experiments were done in a 7T horizontal magnet (Agilent-Technologies) and Bruker electronics (BioSpec AV3HD), using a 35 mm birdcage coil (Doty Scientific). Mice were anesthetized with 2% isoflurane in O2 during data acquisition. T2-weighted axial images were acquired with a TurboRARE-sequence (TR/TE= 2446/42 ms, FOV = 35×35 mm^2^, image size 256×256 and 21 slices of 1.3 mm thickness and non-fat-suppression). Tumor volume analyses were performed by manual segmentation using in-house written scripts in Matlab (Mathworks Inc.).

### Cell lines

Male-derived murine RAW 264.7 macrophages, TRAMP-C2 and TRAMP-C3 prostate cancer cell lines were purchased from ATCC, maintained and cultured according to their suggested protocols. ERG^−^ (PT-83, PT-25, PT-09) and ERG^+^ (PTE-24, PTE-82) syngeneic cell lines (all *Pten*- and *Trb53*-deficient) were generated from tamoxifen treated male mice bearing end-stage prostate tumors as described by others (14). All cell lines lacked expression of CD45 or smooth muscle actin (α-SMA), expressed variable levels of androgen receptor, and expressed detectable keratin 14 and 18. Cells were maintained in Dulbecco modified Eagle’s minimal essential medium (DMEM), 25.0 mM glucose, 25.0 mM HEPES supplemented with 10% fetal calf serum (FCS), 2% penicillin/streptomycin, 0.005 mg/ml bovine insulin and 10 nM dehydroisoandrosterone (DHEA). Subculturing was performed when cells reached 90% confluency and only passage numbers below 10 were used for *in vivo* experiments.

### Primary cells

Bone marrow-derived macrophages (BMDMs) were generated as described previously (15,16). In brief, bone marrow was flushed from femurs and tibias of male C57BL/6J mice and cultured at 37°C for 6-7 days in complete macrophage medium (Dulbecco modified Eagle’s minimal essential medium (DMEM), 25.0 mM glucose, 25.0 mM HEPES, supplemented with 10% fetal calf serum (FCS), 2% penicillin/streptomycin, and 20 ng/ml M-CSF (R&D Systems).

### Flow cytometry

Mice were cardiac perfused with PBS containing 10 U/ml heparin to clear peripheral blood followed by surgical excision of prostate tissue and/or tumors. Single cell suspensions were then prepared by incubating minced tissue in 1 mg/ml collagenase (Roche) and 50 U/ml DNase I (Roche) at 32°C with agitation for 30 min. The resulting cell suspensions was passed through a 70 μm cell strainer and washed once with FBS-containing DMEM followed by once with PBS. Single cell suspensions were then incubated in ACK Lysing Buffer at RT for 3-5 min. Single cell suspensions were then stained immediately or stored in freezing medium (5% charcoal-stripped serum [CSS], 10% DMSO and 85% DMEM) at −80°C for later analysis. Staining protocol involved incubation with live/dead (L/D) fixable fluorescent reactive dyes (Molecular Probes) in PBS for 10 min. After washing, cells were incubated for 30 min at 4°C in 100 μl staining buffer (PBS, 2% BSA) containing antibodies. For intracellular lipid staining, single cell suspensions were incubated with 0.25 μM Bodipy™ 493/503 in PBS for 15 min at RT. Paraformaldehyde Solution, 4% in PBS (Affymetrix) was used to fix cells. Data was recorded on a LSR II Flow Cytometer (BD Biosciences) and analysis completed using FlowJo software.

### LDL uptake

Tumor cells were seeded in culture plates and starved for 48 h in 5% CSS DMEM. RAW 264.7 cells were starved for 24 h in 5% CSS DMEM then labeled with pHrodo™ Red-LDL (L34356) 1:2000 for the last 3 h. Tumor cells were then co-cultured with labeled RAW 264.7 cells for another 24 h in serum-free medium. Samples were then processed for flow cytometry by staining with CD45-APC and DAPI. Cells were mixed just prior to acquisition on the LSRII for the 0 min incubation.

### Co-culture

3000 tumor cells were seeded in each well of 96 well plate, then 1000 macrophages were added on the following day in DMEM (phenol red free, contains 25 mM glucose and 25 mM HEPES) supplemented with charcoal stripped serum (CSS) with or without serial dilutions of enzalutamide (10, 20, 40, 80 μM). Time lapse microscopy measurements were performed every 6-8 h in phase-contrast white light using Incucyte Zoom system (Essen Bioscience). Percent confluency was calculated for each time point using a tailored algorithm for each cell line, and data were normalized to initial time point of each well.

### Confocal immunofluorescence

Cells on chamber slides were washed twice with PBS, fixed with Paraformaldehyde Solution, 4% in PBS for 20 min and permeabilized with 0.1% Triton X-100 for 5 min. Cells were washed twice with PBS, blocked with 2% BSA in PBS for 1 hr and subsequently incubated with anti-androgen receptor antibody (1:400, Abcam) at 4°C overnight. Cells were washed 3 times with PBS and incubated with appropriate fluorescent-labeled secondary antibodies at RT for 1 hr, followed by incubating with F4/80-APC (1:400) for 1h RT in the case of macrophage co-culture experiments. VECTASHIELD^®^ Antifade containing DAPI was used as mounting medium. Vybrant™ Alexa Fluor™ 555 Lipid Raft Labeling Kit was utilized according to the manufacturer’s instructions. Images were visualized using Leica TCS SP8 laser scanning microscope (Leica Microsystems). The nucleus to cytoplasm AR intensity ratio was plotted using the basic (generic) R command hist using R studio version 4.0.3. A smooth curve was fitted to the histogram using kernel density estimation computed by the density function.

### Histology and immunohistochemistry (IHC)

Histological specimens were embedded in paraffin, sectioned (4 μm slices) and left unstained or stained with haematoxylin & eosin (H&E). For immunohistochemistry, slides were stained using a Ventana Discovery XT automated system (Ventana Medical Systems). Briefly, slides were deparaffinized on the automated system with EZ Prep solution (Ventana). Enzymatic retrieval method was used in Protease 1 (Ventana). The rabbit primary antibodies that react to AR (1:200, Abcam), F4/80 (1:100, Abcam), α-SMA (1:250, Abcam), Ly6G (1:100, Biolegend), CD3 (1:200, Abcam), Ki67 (1:300, Abcam), in Dako antibody diluent (Agilent) and incubated for 60 min. The Ventana OmniMap Anti-Rabbit Secondary Antibody was used for 8 min. The detection system used was the Ventana ChromoMap kit, and slides were then counterstained with hematoxylin, followed by dehydration and cover-slipping. Histology slides were scanned using the Aperio™ ScanScope XT with a 200X-(0.8NA) objective lens at a rate of 5 minutes per slide via Basler tri-linear-array. Images were analyzed with Aperio eSlide Manager software using nuclear (AR), membrane (F4/80), or pixel (SMA, Ly6G, Ki67) detection algorithms. Intensity thresholds in the default algorithms for hematoxylin recognition were adjusted to identify faintly stained nuclei. In addition, the positive staining categories (weak, moderate, strong) on the default nuclear detection algorithm were adjusted to account for weaker staining. All Intensity thresholds remained consistent for each image within the respective biomarker group.

### LC-MS/MS Sample Preparation

Stock mixtures of stable isotope-labeled standards were prepared in either methanol or acetonitrile at a concentration of 10 mg/mL and then diluted to lower concentrations. Tumor tissues were collected and stored at −80°C until use. Aliquots of tumor tissue (~50-250 mg) were weighed and placed in Eppendorf microcentrifuge tubes. Samples were kept on ice throughout the processing steps. Each sample was spiked with 10 μL of 9 androgen internal standard mixtures at a concentration of 5 ng/mL for T (2,3,4-^13^C_3_), Prog (2,3,4-^13^C_3_), AED (2,3,4-^13^C_3_) and DOC (2,2,4,6,6,17α,21,21-D_8_), 50 ng/mL (DHT (2,3,4-^13^C_3_), F (9,11,12,12-D_4_) and A5-diol (16,16,17-D_3_), and 500 ng/ml for DHEA (2,2,3,4,4,6-D_6_) and Preg (20,21-^13^C_2_; 16,16-D_2_). After an hour, each sample was re-suspended in PBS, then 0.9-2.0 mm blend stainless steel beads (Next Advance, Troy, NY) were added. The volume ratio of tissue, PBS, and beads is 1:2:1. Samples were homogenized at speed 8 for 5 minutes at 4°C (Bullet Blender, Next Advance, Troy, NY) and transferred to a 5 mL glass centrifuge tube for Liquid-Liquid Extraction (LLE)(17) followed by Solid Phase Extraction (SPE)^(18)^. Briefly, 2 mL of MTBE was added to the glass centrifuge tube with sample homogenate in it and then vortexed for 30 seconds. All samples were centrifuged at 2,000 x g for 10 minutes at 5°C to separate the organic and aqueous layers. The top organic layer was decanted, and the remaining androgens from the aqueous phase were extracted two more times. The pooled MTBE androgen extract was dried under nitrogen (5 psi) using Reacti-Vap™ Evaporators (TS 18826, Thermo Fisher Scientific, Waltham, MA) in a fume hood at room temperature. The androgen pellet was re-dissolved in 2 mL of aqueous 20% methanol and loaded into Isolute C18 cartridges (200 mg/3 mL), which were conditioned with 5 mL methanol and followed by 3 mL water. After loading the sample and washing the cartridges with 2 mL of aqueous 10% methanol, the androgens were eluted with 2 mL 100% methanol. This eluate was dried and reconstituted with 50 μL or 100 μL of aqueous 50% methanol depending on the amount of tissue for each sample. The concentrated extract was transferred to an LC-MS autosampler vial (Fisher Scientific, Hampton, NH) and injected right away.

### LC-MS/MS

Liquid Chromatography-Mass Spectrometry (LC-MS) was performed using a UHPLC (Vanquish, Thermo Scientific) interfaced with a Q Exactive HF hybrid quadrupole-Orbitrap mass spectrometer (Thermo Scientific, San Jose, CA). Chromatographic separation was conducted on an Accucore C18 column (2.1 mm ID x 100 mm in length with 2.6 μm particle size) maintained at 40°C. Separation was achieved using mobile phases A (100% H_2_O with 0.1% formic acid) and B (100% MeOH with 0.1% formic acid). The gradient was programmed as follows: 2 minutes at 30% B, then ramping from 30% B to 90% B over 12 minutes, washing with 90% B for 2 minutes, returning to 30% B over 0.1 minutes and re-equilbrating for 3.9 minutes for a total run time of 20 minutes. The flow rate was set to 0.250 mL/min except 16.1-19 min, where the flow rate was increased to 0.450 mL/min for re-equilibration. For MS, parallel reaction monitoring (PRM) was used for the detection and quantification. The instrument settings were as follows: sheath gas 50, auxiliary gas 10, sweep gas 1, spray voltage 3.5 kV, capillary temperature 325°C, S-lens RF level 50, auxiliary gas temperature 350°C, resolution 30,000, AGC target 2E5, and isolation window width 1.5 m/z with an offset of 0.4 m/z. The data analysis is performed using XCalibur. A processing method was created based on the precursors and fragments listed in **Fig. S2**. The m/z and RT tolerance values were 5 ppm and 0.05 min. The sum of the peak heights of all the fragments for each analyte is used for quantification. To calculate the amounts of androgen in the sample, the ion signal for the endogenous androgen was divided by the ion signal for the corresponding SIS and multiplied by the amount of SIS (in pg) spiked into each sample that was injected on column. To enable more effective comparison between samples, final results are normalized by protein amount and expressed as pg androgen/mg protein.

### RNA sequencing

Mice bearing orthotopic prostate tumors were cardiac perfused with PBS (without heparin), following by resection of tumors and immediate freezing in liquid nitrogen. RNA from prostate tumor tissue was extracted using RNeasy^®^ Plus Mini kit (Qiagen) according to manufacturer’s protocol with on-column DNase digestion and screened for quality on an Agilent Tape-Station (Agilent Technologies, Inc., Wilmington DE). RNA-sequencing libraries were prepared using the NuGen Ovation Mouse RNA-Seq Kit System (Tecan US, Inc., Morrisville, NC). Briefly, 100 ng of DNase treated RNA was used to generate cDNA and a strand-specific library following the manufacturer’s protocol. Library molecules containing ribosomal RNA sequences were depleted using the NuGen Mouse AnyDeplete probe-based enzymatic process. The final libraries were assessed for quality on the Agilent TapeStation (Agilent Technologies, Inc., Wilmington DE), and quantitative RT-PCR for library quantification was performed using the Kapa Library Quantification Kit (Roche Sequencing, Pleasanton, CA). The libraries were sequenced on the Illumina NextSeq 500 v2 sequencer with a 75-base paired-end run in order to generate about 40 million read pairs per sample.

### RNA-seq analysis

The raw reads were first assessed for quality using FastQC (http://www.bioinformatics.babraham.ac.uk/projects/fastqc/) and then trimmed by cutadapt(19) (version 1.8.1) to remove reads with adaptor contaminants and low-quality bases. Read pairs with either end too short (<25bps) were discarded from further analysis. Trimmed and filtered reads were aligned to the GRCm38 mouse reference genome using STAR(20) (version 2.5.3a). Unmapped reads were further aligned to GRCh37 human reference genome. Uniquely mapped reads were counted by htseq-count(21) (version 0.6.1) using Gencode m21 annotation for mouse genome and Gencode v30 annotation for human gene ERG. Differential expression analysis was performed using DESeq2 taking into account of RNA composition bias(22). Genes with fold-change>2 and false discovery rate (FDR) q-values< 0.05 were considered differentially expressed. For each differential expression analysis comparing treated groups vs. IgG group, mouse genes were converted to their human homologs and ranked based on −log10(p-value)*(sign of log2(fold-change)). The ranked gene list used to perform pre-ranked gene set enrichment analysis (GSEA(23) version 4.0.2) to assess enrichment of hallmarks, curated gene sets, and gene ontology(24) terms in MSigDB(23). The resulting normalized enrichment score (NES) and FDR controlled p-values were used to assess the transcriptome changes induced by Lupron and α-CSF1.

### Real-time quantitative PCR

RNA was extracted using RNeasy isolation kit (Qiagen). Real-time quantitative PCR (RT-qPCR) was then carried out using TaqMan™ RNA-to-CT™ 1-Step Kit (Thermo Fisher Scientific). PCR was performed using individual TaqMan Assays with catalogue numbers provided in supplementary table 1 with searchable Research Resource Identifier (RRID). Relative Expression/Fold Change was calculated by using 2^ΔCt^ method where ΔCt =Average Ct Gene of Interest - Average Ct of *Tbp* as Endogenous Control.

### TCGA PRAD analysis

The correlation of macrophage-related genes and de novo cholesterol biosynthesis in a prostate cancer cohorts was computed using level 3 gene expression estimates from the RNA-Sequencing in the TCGA PRAD (Prostate Adenocarcinoma Data Set) database, extracted, and hosted by cBioPortal for Cancer Genomics (www.cbioportal.org).

### GRID registry

To evaluate the clinical relevance of macrophages in therapeutic resistance to ADT we utilized 635 prostatectomy tumor samples with whole transcriptome data from the Decipher GRID^™^ registry data (NCT02609269, institutional review board approved)(25). Gene expression markers of tumor macrophages were used to evaluate their correlation with the ADT response score, a validated gene expression signature used in the prediction of postoperative ADT response, as has been described(26).

### Statistical Analysis

Unless otherwise indicated, unpaired t-test was used to facilitate comparisons between two group means. For multiple group comparisons two-way ANOVA was used. Data points represent biological or technical replicates and are shown as the mean ± SEM or SD as indicated in the figure legends. Intergene correlation matrix with hierarchical clustering, based on spearman correlation coefficient, was used determine correlation clusters. All statistical tests were two sided and are shown as *p<0.05, **p <0.01, ***p<0.001 (described in each figure legend). Statistical analysis was carried out using R studio v3.5.0 and GraphPad PRISM 7 software.

## Results

### Development of syngeneic mouse models of prostate cancer

Prostate cancers often harbor recurrent rearrangements of the E26 Transformation-Specific (ETS) family of transcription factors. Among these, fusion of ETS-related gene (*ERG*) to the membrane protease TMPRSS2 (*TMPRSS2-ERG*) represents the most common structural rearrangement in prostate cancer (27). We established a mouse model that reflects the mutational landscape of aggressive prostate cancer by crossing *Trp53^lox^, Pten^loxP^* and *Pbsn-cre/Esr1*\* mice, with or without the *TMPRSS2/ERG* fusion gene **(Fig. 1A)**. Cre-mediated recombination was confirmed in the mouse prostate 2 months following the initiation of tamoxifen, as shown by immunohistochemistry staining of GFP and increased Ki67 positivity **(Fig. 1B**). Both *Pten*^pce−/−^Trp53^pce−/−^ and *Pten*^pce−/−^Trp53^pce−/−^ *TMPRSS2-ERG*^pce+^ mice began to develop tumors after 6 months, as detected by MRI, with all mice developing tumors within 12 months (**Fig. 1C**). In line with the putative oncogenic role of ETS fusions in prostate cancer(28), there was an increase in tumor incidence and progression, and reduced survival in *TMPRSS2-ERG*^pce+^ mice (**Fig. 1C**). Neoplastic progression at 6 and 12 months (**Fig. 1D**) was marked by higher Ki67 positivity and reduced presence of SMA^+^ fibromuscular stroma in both genotypes (**Fig. S1A-B**). Early tumor lesions were androgen driven as mice (4-month post tamoxifen) treated with Lupron, a gonadotropin-releasing hormone analogue that reduces systemic testosterone levels, displayed decreased tumor lesion size in both genotypes, as shown by representative MRI T2-weighted images **(Fig. 1E, S1C-D).** From end stage mice we generated a series of syngeneic *TMPRSS2-ERG*^+^ (denoted as PTE) and *TMPRSS2-ERG^−^* (denoted as PT) prostate cancer cell lines that expressed varying levels of cytokeratin 8, 14, and 18 (**Fig. 1F**) as well as AR (**Fig. 1G**). As expected, *Pten*-deficiency resulted in sensitivity to the PI3K inhibitor GDC-941 (**Fig. S1E**). AR nuclear translocation was observed following the addition of dihydrotestosterone (DHT) in all cell lines tested (**Fig. 1H**). However, cell lines displayed differential sensitivity to enzalutamide, including 3 lines (PTE-82, PT-09, PT-25) that survived even in the presence of 80 μM enzalutamide, thereby representing models of CRPC (**Fig. 1I**).

**Figure 1.**
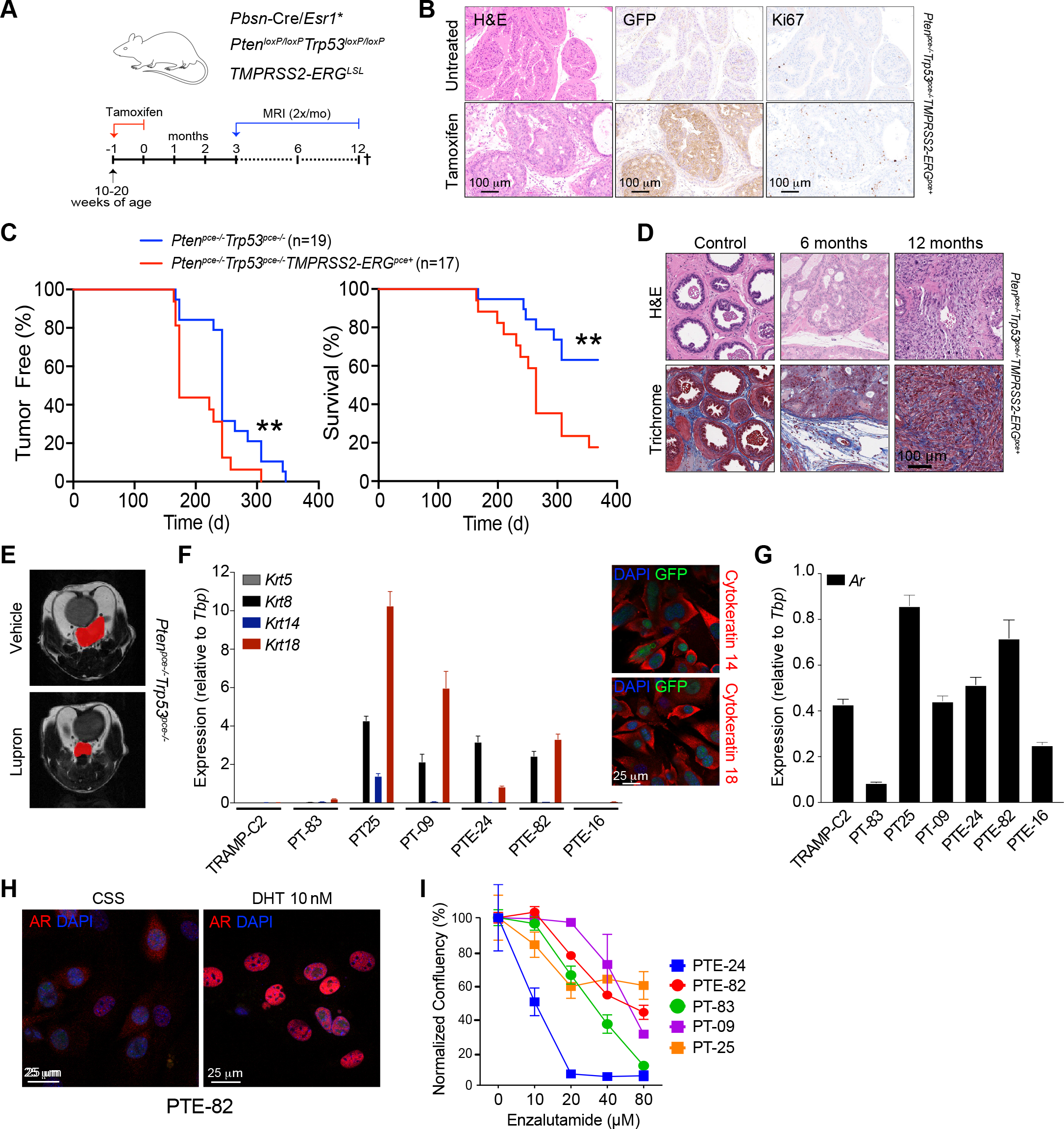
Development of syngeneic mouse models representing CRPC. A) Mouse model scheme. B) Tamoxifen induced GFP and Ki67 expression in prostate glands of a *Pten*^pce−/−^/*Trp53*^pce−/−^ */TMPRSS2-ERG*^pce+^ mouse after 2 month of tamoxifen administration, n=1. C) Detection of prostate tumors and overall survival in *Pten*^pce−/−^/*Trp53*^pce−/−^ and *Pten*^pce−/−^/*Trp53*^pce−/−^/*TMPRSS2-ERG*^pce+^ mice. Days reflect time post tamoxifen administration. Tumor incidence and size were monitored by monthly MRI starting at 5-6 months. n=17-19 mice, pooled from two independent cohorts. Significance between the two groups was determined by log-rank. D) Hematoxylin and eosin (H&E) and Masson’s trichrome staining in *Pten*^pce−/−^/*Trp53*^pce−/−^/*TMPRSS2-ERG*^pce+^ mice from C. Control prostate glands were harvested from mice at 6 and 12 months of age. E) Representative T2-weighted image by MRI of tumor lesion-bearing *Pten*^pce−/−^/*Trp53*^pce−/−^ mice treated with vehicle or 0.6 mg Lupron subcutaneously every 28 days for two cycles, starting 4 months after tamoxifen administration. n=3-5 mice per group, data from a one cohort of mice. F) Expression of *Krt5*, *Krt8*, *Krt14*, *Krt18* in the different cell lines generated from either *Pten*^pce−/−^/*Trp53*^pce−/−^ (PT) or *Pten*^pce−/−^/*Trp53*^pce−/−^/*TMPRSS2-ERG*^pce+^ (PTE) mice at end-stage. n=6 from one experiment, with data shown as the mean ± SEM. TRAMP-C2 cell line was used as a comparison. Representative confocal microscopy immunofluorescent images of the PTE-82 cell line stained for cytokeratin 14 and cytokeratin 18 are shown to the right. Similar results were obtained for the PT-09, PT-25, and PTE-24 lines. Images represent one of two independent experiments. G) Expression of *Ar* in the different cell lines. n=3 from one experiment, data shown as the mean ± SEM. H) Confocal microscopy immunofluorescent images of the PTE-82 cell line either treated with 10 nM DHT or left untreated in charcoal stripped serum (CSS). Androgen receptor (red) and DAPI (blue). Images are representative of one of three independent experiments. Dose response curve of enzalutamide in 5 different prostate cancer cell lines. Phase contrast images were acquired at 8 hr intervals using Incucyte, with confluence per well calculated per cell line. n=3, data shown as the mean ± SEM from one of at least three independent experiments.

### Macrophages are associated with cholesterol transport and androgen synthesis

Both the PT-09 and PTE-82 CRPC cell lines established orthotopic tumors following injection into the prostate glands of C57BL6/J mice. Critically, F4/80^+^ macrophages formed the dominant immune subset in these orthotopic tumors, with Ly6G^+^ neutrophils comprising only a small percentage of the total CD45^+^ population (**Fig. 2A**). This recapitulated our findings in autochthonous *TMPRSS2-ERG*^pce+^ and *TMPRSS2-ERG*^pce-^ prostate tumors at both 6 and 12 months (**Fig. S1A-S1B**). In solid tumors the majority of TAMs differentiate from peripheral blood monocytes under the influence of colony stimulating factor-1 (CSF-1), with a smaller population of tissue-resident macrophages that persist through in situ proliferation (29–31). To determine the role of macrophages in CRPC, we treated mice bearing orthotopic PTE-82 tumors with an αCSF-1 neutralizing antibody for 7 days **(Fig. 2B)**, which reduced the percentage of CD11b^+^F480^+^ TAMs without altering the frequency of other immune subsets (**Fig. S2A**). Another group of mice were treated with 0.6 mg/mouse Lupron Depot 16 days prior to tumor collection and processing for RNA sequencing to provide a comparison to standard-of-care therapeutics.

**Figure 2.**
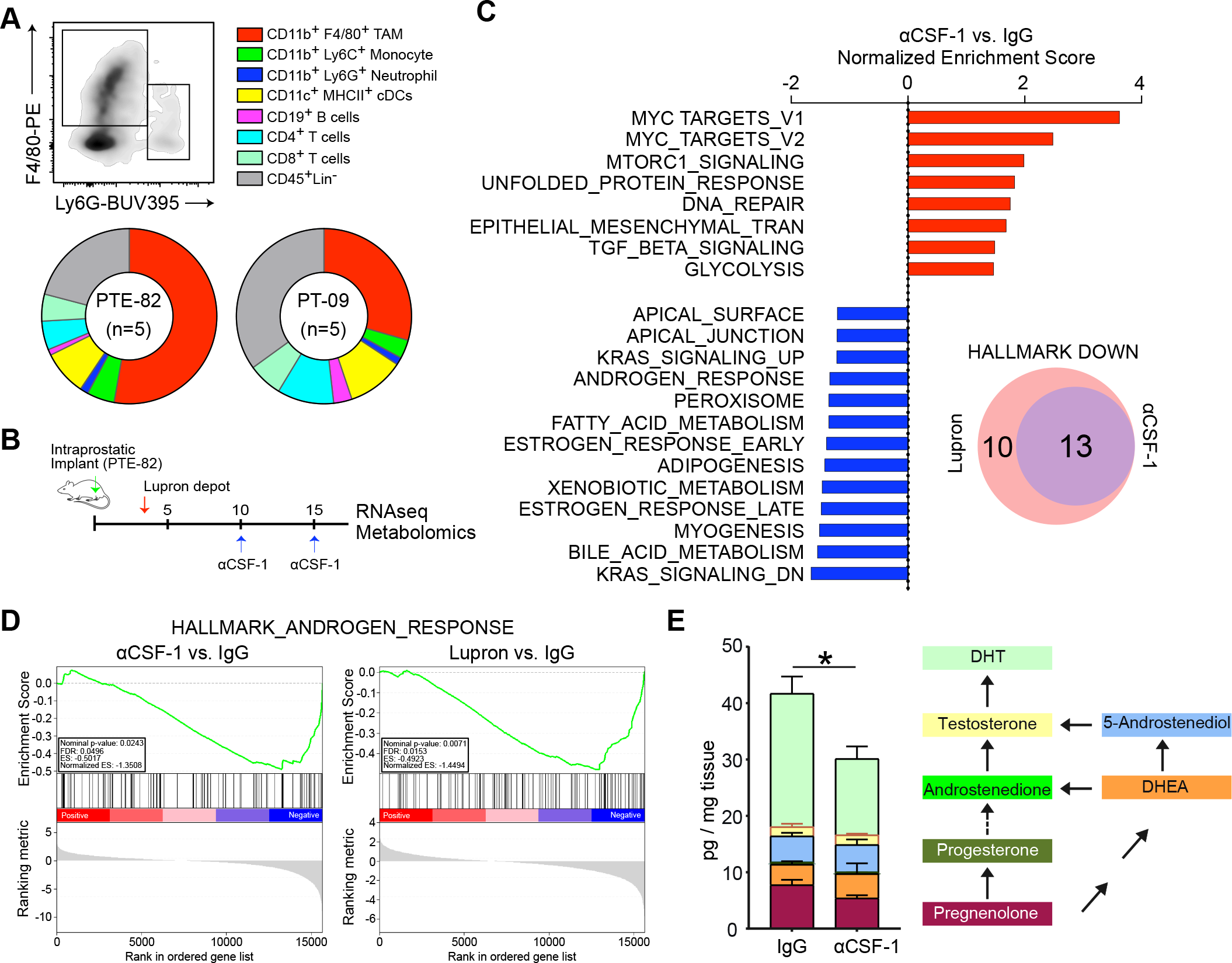
Macrophages are associated with cholesterol transport and steroid hallmarks. A) Immune infiltration in orthotopic models of prostate cancer, as determined by flow cytometry. n=5 mice per group, with results from one of three independent experiments shown as the mean. B) Treatment schematic for Lupron and α-CSF1. C) Gene set enrichment analysis (GSEA) following whole tumor RNA sequencing. The normalized enrichment score is shown comparing tumors from mice treated with α-CSF1 versus an IgG control. Data reflects 3 tumors per group from a single experiment. D) Enrichment plots of the Androgen Response Hallmark show for tumors from mice treated with α-CSF1 or Lupron, compared to the IgG control. E) LC-MS/MS quantification of pregnenolone, progesterone, androstenedione, DHEA, 5-androstenediol, testosterone and DHT in fresh tumor tissues. n=5-6 mice per group, from one of two independent experiments. Measurements were normalized by tissue weight. Significance determined by unpaired t-test for DHT. *p<0.05.

As shown in **Fig. 2C,** RNA sequencing followed by gene set enrichment analysis (GSEA) revealed that αCSF-1 treatment decreased the expression of genes related to steroid metabolism, bile acid metabolism and adipogenesis. Interestingly, these changes were very similar to those observed following treatment with Lupron (**Fig. 2C, S2B**). Depletion of TAMs elicited decreased expression of many AR-regulated genes (e.g. endogenous *Tmprss2*), along with a decrease in the global AR signature (normalized ES −1.3508) nearly equivalent to Lupron (normalized ES −1.4494) (**Fig. 2D-S2C**). As cholesterol is the main precursor of bile acids and the five major classes of steroid hormones, the results suggested that depletion of TAMs was causing a perturbation in cholesterol transport and/or metabolism. We reasoned that this perturbation could interfere with androgen biosynthesis in the tumor microenvironment, thereby explaining the reduced AR signature observed following TAM depletion. To evaluate this possibility, we utilized a LC-MS/MS metabolomics platform to quantify the level of intra-tumoral androgens in mouse tissues (**Fig. S2D**) (32). As shown in **Fig. 2E**, αCSF-1 treatment decreased the overall level of steroids, including the most potent endogenous androgen, DHT. Combined, these data indicate that macrophages regulate androgen biosynthesis in CRPC, potentially through a role in cholesterol transport/metabolism.

### Macrophages directly regulate AR nuclear translocation and resistance to enzalutamide

The AR functions by binding steroid hormones in the cytoplasm and translocating to the nucleus, where, in conjugation with co-factors, it regulates expression of target genes by binding to specific DNA sequences known as androgen response elements (33). Based upon the reduction in intratumoral DHT and the AR gene signature that followed depletion of TAMs (**Fig. 2D-E**), we measured the amount of AR nuclear localization within tumors following administration of αCSF-1. Nuclear AR staining was prevalent in control tumors (**Fig. 3A**) and correlated with infiltration by F4/80^+^ macrophages (**Fig. S3A**). In contrast, the reduction in TAMs that followed administration of αCSF-1 coincided with a significant reduction in nuclear AR (**Fig. 3A-B**). This was not due to reduced AR protein expression in αCSF-1 treated tumors, but rather sequestration of AR in the cytosol. Notably, the reduction in AR nuclear translocation following TAM depletion was equivalent to treating mice with Lupron (**Fig. 3A**). In addition, upon analyzing autochthonous *TMPRSS2-ERG*^pce+^ at 6 and 12 months, we found a positive correlation between nuclear localization of AR and the extent of macrophage infiltration (**Fig. S3B**). Intriguingly, this correlation was not observed in autochthonous *TMPRSS2-ERG*^pce-^ tumors (**Fig. S3B**). Thus, TAMs regulate AR activation and nuclear translocation, at least in CRPC tumors expressing an ETS fusion gene.

**Figure 3.**
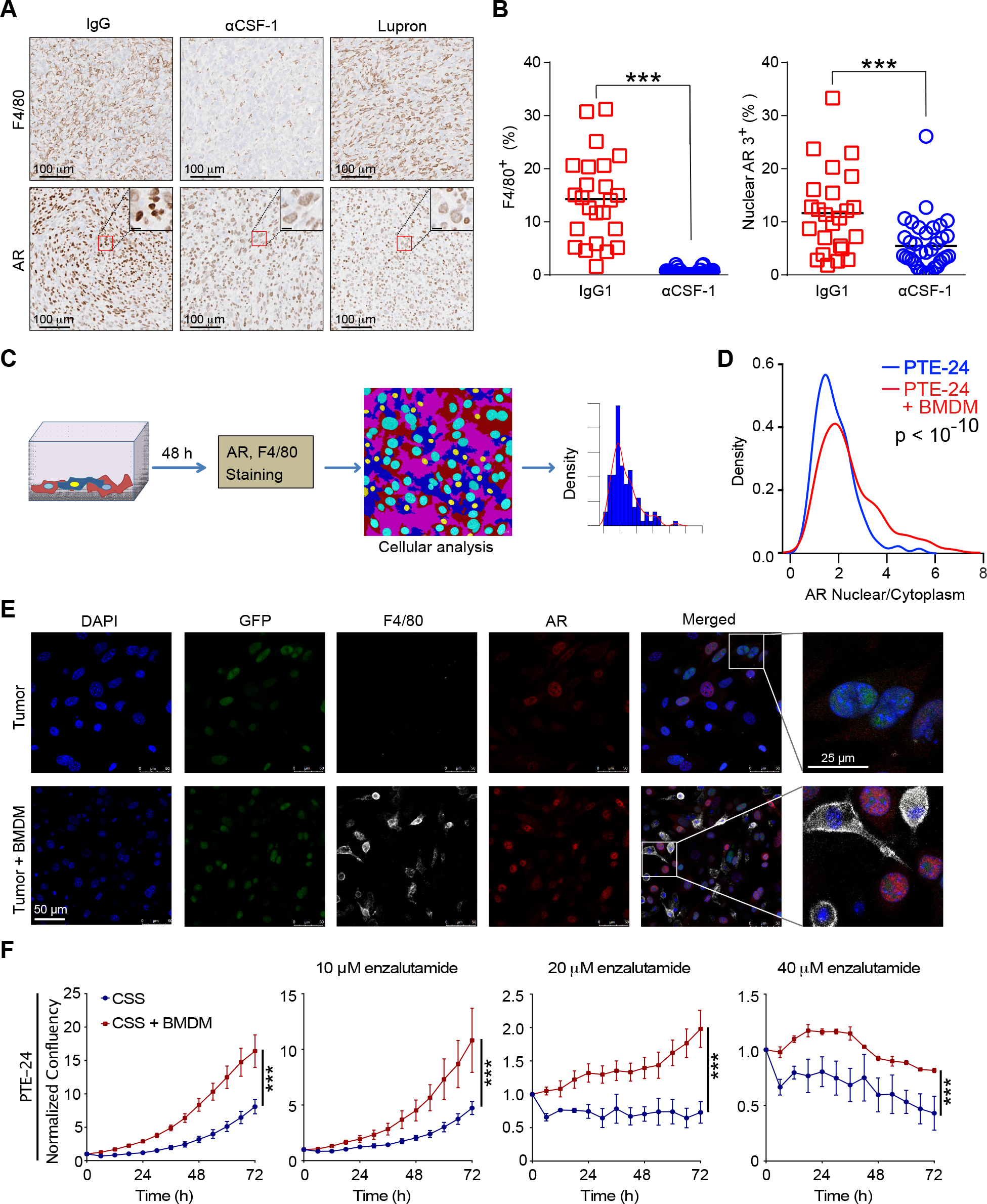
Macrophages directly regulate AR nuclear translocation. A) Representative immunohistochemistry for tumor macrophages (F4/80) and AR within orthotopic prostate tumors from mice treated with IgG (control), α-CSF or Lupron. B) Quantification of the percent of F4/80^+^ cells and AR^+^ nuclei in serial sections. n=5-6 mice per group from one experiment. 4-5 regions of interest (ROI) per each slide were selected in non-necrotic areas of the tumor. Significance determined by unpaired t-test and shown as ***p<0.001. C) Experimental schematic for cancer cell and BMDM co-culture and quantification of nuclear AR by confocal microscopy. D) Kernel density estimation of the AR nuclear to cytoplasmic ratio on a cell-by-cell basis following incubation of PTE-24 cancer cell line alone, or in co-culture with BMDMs under androgen-deprived conditions (i.e., charcoal-stripped serum, CSS) for 72 hrs. n=3, data from one of at least three independent experiments. Significance determined by Mann-Whitney. E) Representative confocal microscopy images of the GFP^+^ PTE-24 cell line. AR (red), F4/80 (white) and DAPI (blue) fluorescence is shown. F) Impact of BMDMs on the proliferation of PTE-24 cells in the presence of serial concentrations of enzalutamide. Cell proliferation was monitored using live imaging with phase contrast images acquired every 6 hr. Data shown as the mean ± SEM and reflects one of 2 independent experiments. Significance determined by two-way ANOVA.

We next sought to determine whether macrophages could directly induce AR nuclear translocation. We first established an *in vitro* system using a castration-sensitive cell line that was highly dependent upon AR signaling for growth, PTE-24 (**Fig. 1I**). PTE-24 cells were co-cultured with bone marrow-derived macrophages (BMDMs) under androgen-deprived conditions (i.e., 5% CSS) and AR nuclear localization was measured via confocal microscopy and quantified by the ratio of nuclear to cytoplasmic AR on a cell-by-cell basis (**Fig. 3C**). As shown in **Fig. 3D-E**, the addition of BMDMs induced a significant increase in AR nuclear localization. As a result of this androgen-induced AR activation, BMDMs promoted the proliferation of PTE-24 cells in androgen-deprived conditions and protected against the cytotoxicity of the AR inhibitor enzalutamide (**Fig. 3F**). AR nuclear localization was also enhanced during co-culture of BMDMs with the PT-09 and PTE-82 CRPC cell lines (**Fig. S3C**); however, these lines are resistant to enzalutamide (**Fig. 1I**) and coculture with BMDMs had only a minor impact on proliferation by PTE-82 cells (**Fig. S3D**).

### Macrophages regulate cholesterol bioavailability

We were intrigued by our comparative transcriptomic studies showing downregulation of bile acid- and steroid biosynthesis-related genes in tumors isolated from αCSF1 treated mice, which suggested altered cholesterol metabolism. We thus performed correlation analysis between macrophage-related genes (*CSF1R*, *CD68*, *CD206*) and genes involved in bile acid and lipid metabolism in TCGA data (Prostate Adenocarcinoma dataset, Cell 2015)(34). We observed that macrophage-related genes correlated with those related to cholesterol transport, such as *ABCA1*, *ABCA8*, *ABCA6* (**Fig. 4A**). Since cellular cholesterol is maintained by a balance between uptake and synthesis, we queried whether *de novo* cholesterol synthesis might be reduced in the presence of high macrophage infiltrate, and observed an inverse correlation between macrophage markers and *de novo* cholesterol synthesis-related genes (*ACAT2*, *HMGCR*, *HMGCS1*, *SQLE*) in two prostate cancer patient datasets (**Fig. 4B, S4A**).

**Figure 4.**
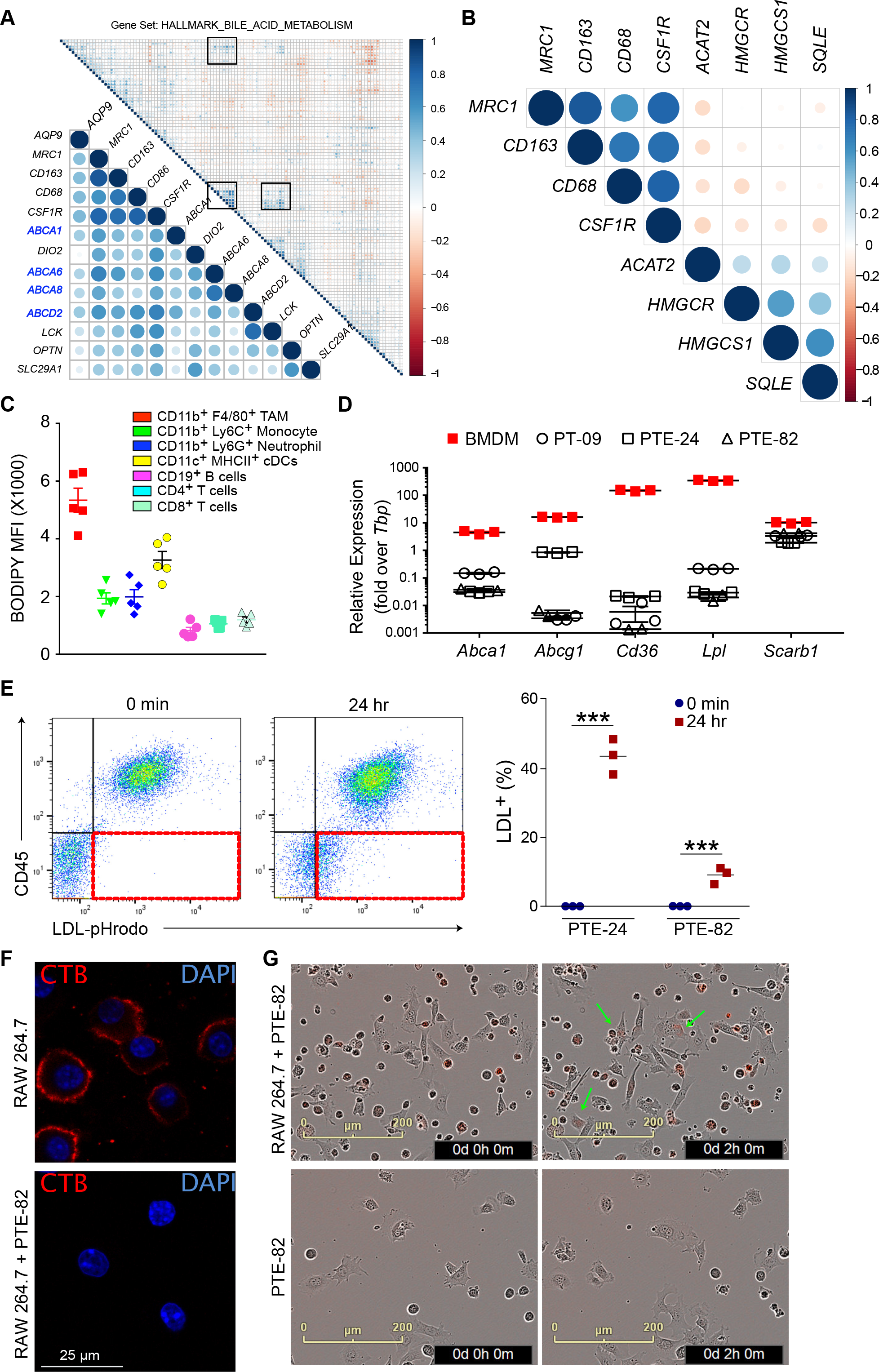
Macrophages transfer cholesterol to prostate cancer cells. A) Intergene correlation analysis of genes associated with macrophage infiltration (*CD86*, *CSF1R*, *MRC1*, *CD163*) and bile acid metabolism from TCGA. B) Intergene correlation analysis of genes associated with macrophage infiltration and *de novo* cholesterol synthesis (*ACAT1/2*, *HMGCR*, *HMGCS1*, *SQLE*). Analysis was performed with the Corrplot package in R. Correlation coefficients are indicated by changes in circle color and size. C) Neutral lipid staining in immune cells within orthotopic PTE-82 tumors, as measured by BODIPY-493/503 staining in single cell suspensions. n=5 mice, data shown as the mean ± SEM from one of three independent experiments. D) Expression of cholesterol metabolism-related genes in prostate cancer cell lines and BMDMs. n=3, data from one of three independent experiments. E) RAW264.7 cells were loaded with human LDL covalently conjugated to pHrodo Red for 3 hrs, then co-cultured with prostate cancer cells for 24 hrs prior to flow cytometric analysis. Cells were cultured separately and then admixed just prior to data acquisition for the 0 min control. The acquisition of fluorescence by the CD45^−^ cancer cells is highlighted in the 3^rd^ quadrant in red. Quantification of percent LDL-positive tumor cells is shown to the right. n=3 biological replicates from one of two independent experiments. Data are presented as mean ± SEM. Student’s *t* test was utilized for statistical analysis and is shown as \*\*\**p*<0.001. F) Immunofluorescence staining of cholesterol rich lipid rafts in the plasma membrane of RAW264.7 cells using Cholera toxin B (red). Cells were cultured alone or with PTE-82 cells in CSS for 48 hrs. Only nuclear GFP^−^ RAW264.7 cells are shown. n=3, data from one of two independent experiments. G) Live imaging of uptake of pHrodo Red-LDL (0.5 μg/ml) by PTE-82 tumor cells in the presence or absence of RAW264.7 cells. Arrows indicate tumor cells with red fluorescence. n=3, data reflects one of three independent experiments.

Cholesterol is mostly stored in the form of cholesteryl esters in intracellular lipid droplets. Accordingly, we quantified the lipid content of immune cells within tumors and found that TAMs had the highest level of lipid content, both in orthotopic and autochthonous *TMPRSS2-ERG*^pce+^ tumors (**Fig. 4C, S4B-C**). BMDMs also expressed numerous genes related to cholesterol influx/efflux at higher levels than the prostate cancer cell lines, including *Abca1, Abcg1, Cd36, Lpl* and *Scarb1* (**Fig. 4D**). To evaluate if macrophages were capable of transferring cholesterol to tumor cells, we loaded RAW264.7 cells, a murine macrophage cell line, with pHrodo-Red conjugated LDL as a source of cholesterol (and associated fatty acids and proteins). Note that pHrodo is fluorescent only at an acidic pH, signifying intracellular uptake via endocytosis and/or phagocytosis has occurred (**Fig. 4E**). Preloaded RAW264.7 cells were then co-cultured with the prostate cancer cell lines for 24 hours, and depending on the tumor cell line, up to 40% of the cancer cells acquired derivate components of LDL (**Fig. 4E**).

To evaluate whether prostate cancer cells were promoting cholesterol efflux by macrophages, we labeled cells with cholera toxin subunit B (CTB), which is capable of binding to cholesterol-rich lipid rafts in the plasma membrane. As shown in **Fig. 4F**, co-culture with cancer cells largely eliminated CTB labeling of RAW264.7 cells, demonstrating a reduction in plasma membrane cholesterol. Finally, we sought to determine if the presence of macrophages could promote the acquisition of cholesterol (or other LDL components) by adding pHrodo-Red conjugated LDL to either prostate cancer cells alone, or those in co-culture with RAW264.7 cells (**Fig. 4G**). RAW264.7 cells rapidly took up LDL and were mostly positive during the first image acquisition, but fluorescent signal began to appear in the adherent prostate cancer cells within 2 hours. In contrast, no fluorescent signal was observed when prostate cancer cells were incubated alone (**Fig. 4G**). Overall, these data suggest that macrophages are a rich source of cholesterol, are capable of transferring cholesterol to prostate tumor cells, and can increase cholesterol bioavailability.

### Activating LXRβ inhibits macrophage-mediated AR translocation in tumor cells

Our cancer cell lines expressed detectable, albeit low, levels of the first two enzymes required for the conversion of cholesterol into androgens (*Cyp11a*, *Cyp17a*), whereas we were unable to detect gene expression of the enzyme responsible for cholesterol side-chain cleavage (*Cyp11a*) in BMDMs (**Fig. S5A**). We therefore sought to determine if preventing cholesterol transfer between macrophages and tumor cells could restrict AR nuclear translocation as a surrogate of androgen production. MetaCore analysis of our RNAseq data revealed that tumors with high macrophage infiltration (i.e. IgG control) were enriched in a Liver X Receptor (LXR)-regulated transcriptional program including target genes such as *Abca1*, *Abcg1*, *Scarb1 and Cd36*. Macrophages are generally thought to unload cholesterol via efflux mediated via the ATP-binding cassette transporters A1 and G1 (ABCA1 and ABCG1) to apolipoprotein A1 and high-density lipoproteins (HDLs), respectively, a process known to regulate atherosclerosis (35–37). Macrophage expression of these transporters has also recently been shown to promote the growth of ovarian cancer (38). We therefore crossed *Abca1^fl/fl^*/*Abcg1^fl/fl^* mice with *LysM-Cre*^+^ mice to generate myeloid-specific loss of ABCA1 and ABCG1. Surprisingly, the extent of AR nuclear positivity within orthotopic tumors was equivalent in *LysM*-cre^−^ and *LysM*-cre^+^ animals (**Fig. 5A**). There was also no difference in the growth of orthotopic tumors between the two genotypes (**Fig. 5B**). Consistent with this, BMDMs generated from *LysM*-Cre^−^ and *LysM*-Cre^+^ mice were equally capable of inducing AR nuclear translocation (**Fig. 5C**). AR nuclear translocation *in vitro* was similarly unchanged by a blocking antibody against CD36 or the addition of a specific scavenger receptor class B type I (SR-B1, encoded by *Scarb1*) inhibitor, BLT-1 (**Fig. S5B**).

**Figure 5.**
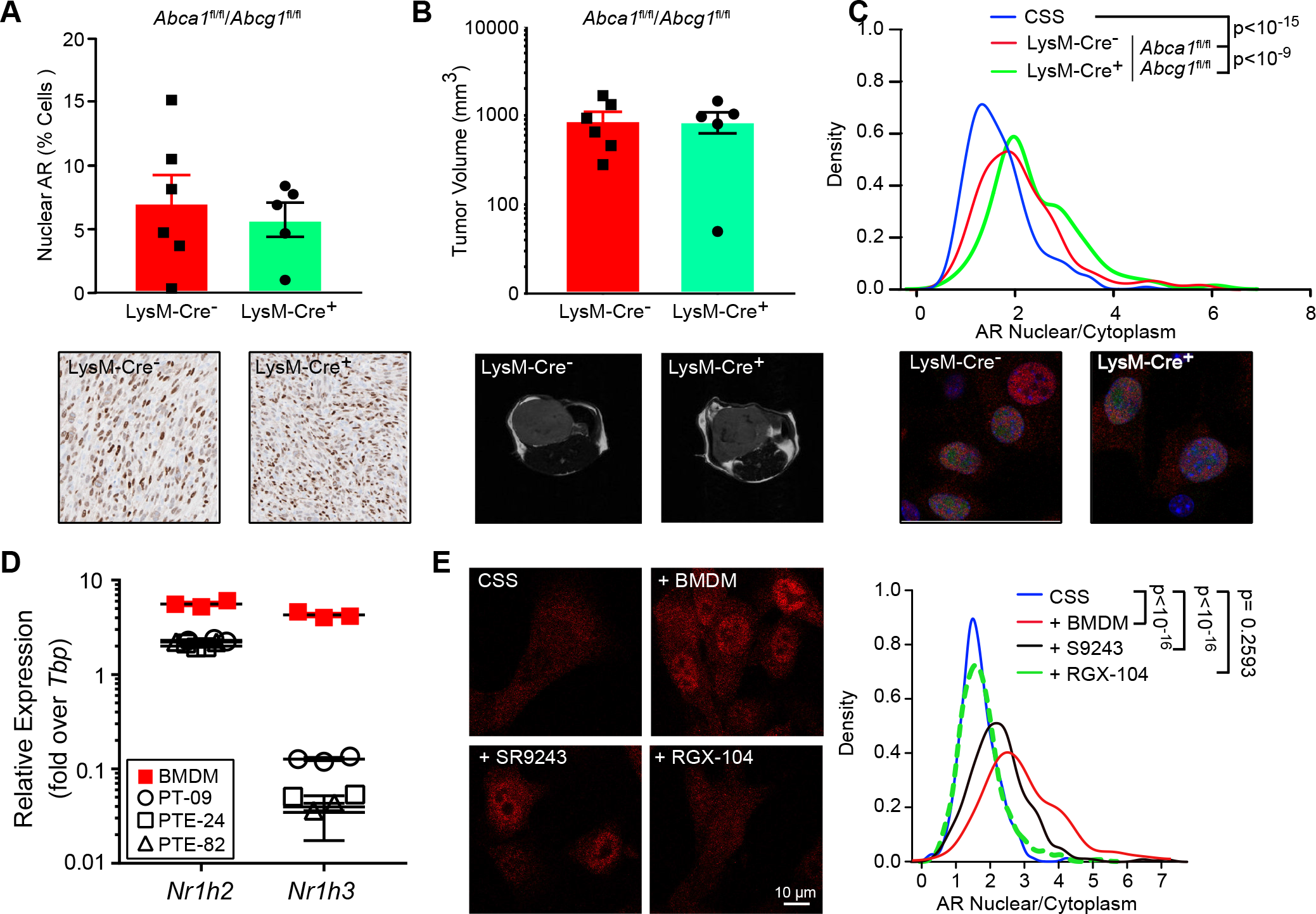
Perturbing cholesterol homeostasis inhibits macrophage-mediated AR translocation. A) PTE-82 prostate cancer cells were orthotopically injected into *LysM*-Cre^+^/*Abca1*^fl/fl^/*Abcg1*^fl/fl^ or *LysM*-Cre^−^/*Abca1*^*fl*/fl^/*Abcg1*^fl/fl^ and nuclear expression of AR was evaluated after 19 days by IHC. Representative images are shown below. n=5-6 mice per group, data shown as the mean ± SEM. Significance was determined by unpaired *t* test assuming Gaussian distribution and with Welch’s correction. B) Tumor volume in mice from A, as measured by MRI. Representative T2-weighted MRI images are shown. C) Nuclear to cytoplasmic AR ratio of GFP^+^ PTE-82 cells cultured in CSS, either alone or in the presence of BMDMs derived from *LysM*-Cre^+^/*Abca1*^fl/fl^/*Abcg1*^fl/fl^ or *LysM*-Cre^−^/*Abca1*^*fl*/fl^/*Abcg1*^fl/fl^ donors. 6-8 randomly selected images from two wells of the chamber slides were pooled for analysis. Data reflects one of three independent experiments. Significance was determined by Mann-Whitney. D) Expression of *Nr1h2* (LXRβ) and *Nr1h3* (LXRα) in prostate cancer cell lines and BMDMs. n=3, data shown as the mean ± SEM from one of two independent experiments. E) Nuclear to cytoplasmic AR ratio of GFP^+^ PTE-82 cells cultured in CSS or co-cultured with BMDMs. 5 μM SR9243 (LXRα/β inverse agonist) or 5 μM RGX-104 (LXRβ agonist) were added to co-cultures as indicated. 6-8 randomly selected images from two wells of the chamber slides were pooled for analysis. Data reflects one of three independent experiments. Significance determined by Mann-Whitney.

It has recently been shown that macrophages have the ability to transfer cholesterol through direct transcellular cholesterol movement in an ABCA1/G1-independent manner (39). Although the molecules involved in this process are unknown, upon re-examining our live imaging data we noticed that pHrodo-Red fluorescence in cancer cells occurred within those in proximity to RAW264.7 cells (**Fig. 4E**). We therefore sought to determine if inhibiting cholesterol influx and/or inducing cholesterol efflux in tumor cells would restrict AR nuclear translocation by activating the LXR transcription factors, which function as the master regulators of cholesterol homeostasis. Both BMDMs and prostate cancer cells expressed detectable levels of the genes encoding LXRα (*Nr1h3*) and LXRβ (*Nr1h2);* however, *Nr1h2* was expressed at much higher levels than *Nr1h3* in the prostate cancer cell lines (**Fig. 5D**). As expected, the LXRαβ inverse agonist, SR943, downregulated expression of *Abca1* in cancer cells, while an LXRβ agonist currently in clinical trials, RGX-104, had the reverse effect (**Fig. S5C**). RGX-104 was also able to functionally inhibit prostate cancer cells from taking up pHrodo-Red LDL during a 24-hour incubation (**Fig. S5D**). We therefore examined the ability of SR943 and RGX-104 to inhibit AR nuclear translocation during BMDM and prostate cancer cell co-culture. As shown in **Fig. 5E**, SR943 only slightly reduced the extent of AR in the nucleus, while RGX-104 completely prevented AR nuclear translocation induced by the presence of BMDMs. This was not due to a direct effect on basal AR activity as expression of the AR target gene, Fkbp5, was not impacted by either the inverse agonist or agonist when BMDMs were not present (**Fig. S5E**). Collectively, these data demonstrate that cholesterol exchange between macrophages and prostate tumor cells enhances AR activation.

### Macrophage depletion sensitizes CRPC to Lupron

Based upon the ability of TAMs to regulate the level of intratumoral androgens (**Fig. 2E**), AR nuclear localization (**Fig. 3A**) and AR-dependent gene expression (**Fig. 2D**), we evaluated the clinical relevance of these findings within the Decipher GRID™ registry data of 635 prostatectomy tumor samples with whole transcriptome data. As macrophages and AR nuclear localization were only correlated in the *TMPRSS2*-*ERG*^pce+^ murine model (**Fig. S3A**) we stratified patients by ETS status and compared the ADT response signature with expression of macrophage-associated genes (*CD68*, *CD163, CSF1R*, or *MRC1*). In ETS^+^ tumors we found that the genomic ADT response signature was inversely correlated with the presence of macrophages, whereas this association was not observed in ETS^−^ tumors (**Fig. 6A**). Accordingly, we evaluated if depleting TAMs could reduce tumor growth and/or enhance survival in the TMPRSS2-ERG^+^ orthotopic CRPC model system (**Fig. 6B**). Treatment with either αCSF1 or Lupron alone had only a small impact on the growth of tumors as measured by MRI; however, a more significant reduction in tumor growth was observed when these therapies were combined (**Fig. 6C, S6A-B**). We therefore extended these studies to evaluate survival as measured by tumor size (<1200 mm^3^), organ obstruction, or overall animal health. Over the course of three independent experiments we observed a significant improvement in survival in mice treated with αCSF1 and Lupron, but not in mice treated with either agent along (**Fig. 6D**). Thus, targeting TAMs can improve response to ADT in ETS^+^ CRPC.

**Figure 6.**
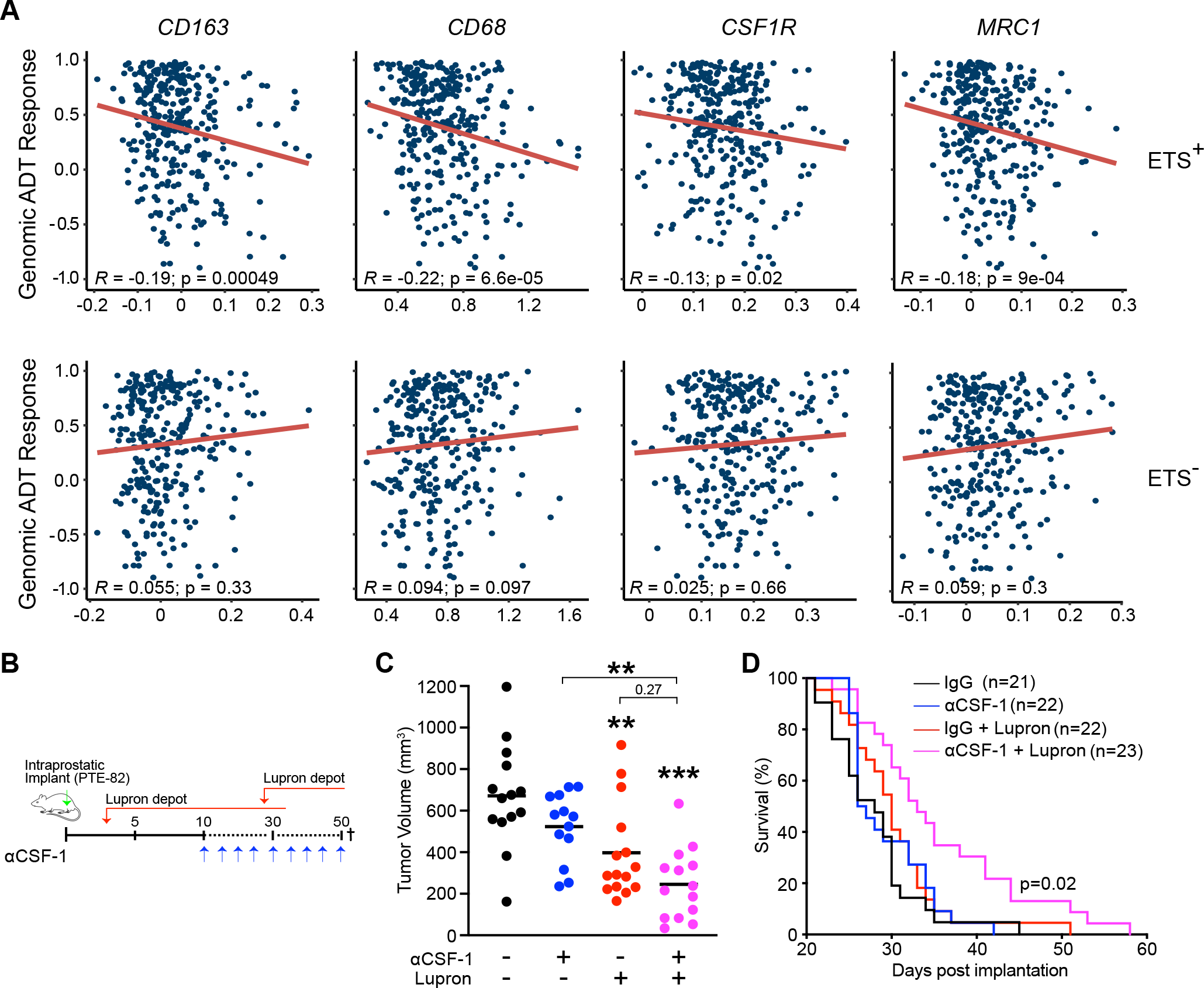
Macrophage depletion sensitizes CRPC to Lupron. A) Expression of macrophage-associated genes was correlated with ADT response score, with stratification by ETS status. B) PTE-82 tumor cells were implanted orthotopically into prostates of C57 mice, which were then treated with Lupron depot and/or αCSF-1 as indicated. Tumor volumes for individual mice are shown for day 21. n=13-15 mice per group, data pooled from two independent experiments. Significance was determined by one-way ANOVA and is show compared to the IgG control group, unless otherwise indicated. C) Tumor volume in treatment groups as measured by MRI. D) Survival as determine by tumor volume or organ obstruction. n=21-23 mice, data pooled from three experiments. Significance was determined via log-rank.

## Discussion

AR signaling remains active in CRPC and plays a critical role in disease progression, as evidenced by the success of enzalutamide and abiraterone in extending survival in the metastatic setting (6). Compensatory growth factors secreted by other cell types in the tumor microenvironment are also thought to promote castration resistance (40). Castrating tumor-bearing mice elicits an influx of leukocytes, including B cells, neutrophils, and macrophages into prostate tumors, which are able to drive the emergence of CRPC by regulating senescence and proliferation (41–45). These studies point towards a feed-forward signaling loop in which cell stress induces expression of chemokines that promote leukocyte accumulation and drive the growth and development of CRPC. Accordingly, reducing infiltration of macrophage and neutrophils through the use of a CSF1R or CXCR2 antagonist, respectively, enhances the efficacy of ADT (42,44). Critically, increased macrophage infiltration has been described in radical prostatectomy samples from ADT-treated patients, and high macrophage infiltration is associated with increased risk of biochemical recurrence (46).

Despite the association between macrophages and CRPC, the mechanism by which TAMs regulate response to ADT is unclear. CD11b^+^Ly6G^+^ myeloid cells in prostate tumors can act as an important source of IL-1 receptor antagonist and IL-23 (44,46), whereas no specific macrophage-derived factors have been shown to drive the development of CRPC. Here we describe how macrophages regulate local steroid biosynthesis via enhancing the uptake of extrinsic cholesterol by prostate cancer cells, thereby leading to AR signaling and resistance to lupron in a model of CRPC. *De novo* steroidogenesis begins with cholesterol translocation into the mitochondria where the rate-limiting enzyme CYP11A1 converts cholesterol to pregnenolone, followed by series of enzymatic reactions to produce various steroids such as androgens, estrogens or glucocorticoids (47). Recently, glucocorticoids produced by TAMs were shown to elicit T cell dysfunction and resistance to immunotherapy in the MC38 colon carcinoma model (11). However, another recent study reported little to no expression of *Cyp11a1* in TAMs within B16-F10 melanoma tumors, instead noting high expression by T cells and granulocytes (48). In our study, we did not detect *Cyp11a1* in BMDMs, pointing towards the transfer of cholesterol rather than pregnenolone or other downstream derivatives *in vitro*. Prostate cancer cells also express most steroidogenic enzymes and have the ability to convert cholesterol to androgens (49–51). However it remains possible that there are other mechanisms by which macrophages regulate steroidogenesis, as reported in early studies investigating modulation of Leydig cell steroidogenesis by testicular macrophages (52).

Lipid transporters and scavenger receptors have emerged as potential targets in various types of solid malignancies (53,54). In the ID8 ovarian cancer model, genetic ablation of *Abca1*/*Abcg1* in myeloid cells decreased tumor growth, although whether this was through altered macrophage function or reduced cholesterol transfer to tumor cells is unclear (38). Similar to this study, we saw that the presence of tumor cells induced cholesterol efflux in macrophages. However, using the same genetic system we did not observe changes in prostate tumor growth or the ability of macrophages to promote AR nuclear localization *in vitro* or *in vivo*. Indeed, we found that macrophages can directly transfer cholesterol in serum free medium lacking lipoproteins, which would limit the ability of ABCA1/G1 to transfer cholesterol onto HDL particles. Inhibition of the HDL receptor SR-B1 also had minimal impact on the ability of BMDMs to promote AR nuclear localization *in vitro*. Our results are therefore in line with a recent a study that reported transcellular transfer of macrophage cholesterol to smooth muscle cells in the absence of serum or HDL (55). Additional work is required to delineate the molecular mechanism by which this transfer occurs.

In addition to *PTEN* and *TP53* alterations, approximately 40% of prostate cancers carry recurrent rearrangements of the ETS family of transcription factors, of which fusion of *ERG* to the membrane protease *TMPRSS2* represents the most common (56). Murine studies have shown that the *TMPRSS2-ERG* fusion increases early onset invasive prostate cancer through changes in the AR cistrome and enhancing AR output (28). *TMPRSS2-ERG* gene fusion status is also associated with a high androgen-regulated gene expression in early onset prostate cancer (57,58). Apart of the transcriptional impact, *TMPRSS2-ERG*^+^ patients have androgen profiles that differ substantially from *TMPRSS2-ERG*^−^ patients, including increased DHT/testosterone ratios (59). These data are consistent with our preclinical observations regarding the correlation between macrophages and AR nuclear translocation in the autochthonous and orthotopic *TMPRSS2-ERG*^+^ murine model, and with the inverse correlation between macrophage infiltration and the genomic ADT response signature in ETS^+^ patients. Cumulatively, these results suggest that macrophage-targeted agents such as CSF1R inhibitors should be directed towards patients with ETS^+^ prostate cancer.

Collectively, our data show that macrophages deliver cholesterol to prostate tumor cells as an example of metabolic cooperation, thereby increasing intratumoral androgen production and activation of the AR. Whether targeting macrophages, cholesterol transfer, or cholesterol metabolism will prove therapeutically viable in prostate cancer remains to be determined. Interestingly, LXRβ agonists induce apoptosis of immunosuppressive myeloid cells through enhanced expression of ApoE, thereby driving T cell responses in melanoma (60–62). Combined with their ability to restrict the cholesterol pool required for tumoral androgen synthesis (51), LXRβ agonists could therefore have profound impacts on the tumor microenvironment and offer unique opportunities for combinatorial therapies. Regardless of the approach taken, it is likely that macrophage-targeted therapies will display limited efficacy as single agents and that combinatorial approaches utilizing ADT should be evaluated clinically.

## Acknowledgements

The authors would like to thank Noel Clark, Sean Yoder, Antonio Ortiz, Jayden Cline and Joseph Johnson for technical assistance. The authors also thank John Cleveland, Conor Lynch, Jingsong Zhang and Paulo Rodriguez for scientific discussion.

## Author Contributions

Conceptualization, A.E. and B.R.; Methodology, A.E. and M.L.; Formal Analysis, A.E., S.A., J.D., X.Y.; Investigation, A.E., W.D.V., M.L., A.K., K.S., L.N., Writing – Original Draft, A.E. and B.R.; Writing – Review & Editing, A.E., K.Y., X.Y., J.K., R.A.G, and B.R.; Visualization, A.E. and B.R.; Supervision, R.G., K.Y., J.K., R.A.G., and B.R; Funding Acquisition, R.A.G. and B.R.

