## Supplemental Information for "Macrophage-derived cholesterol contributes to therapeutic resistance in prostate cancer"

### Supplementary Figures

**Supplementary Figure 1, Related to Figure 1.** A) Representative images from formalin-fixed, paraffin-embedded prostate tissue, isolated from control (age-matched C57BL6/J or untreated transgenic animals) or 2, 6, or 12 months post tamoxifen administration to *Pten*$^{pce-/-}$/*Trp53*$^{pce-/-}$ or *Pten*$^{pce-/-}$/*Trp53*$^{pce-/-}$/*TMPRSS2-ERG*$^{pce+}$ animals. B) Quantification of F4/80 positive cells, or percent pixel positivity for Ly6G, Ki67, and α-SMA, in the tissue from A. n=2-5 age-matched control animals or n=13-16 tamoxifen-treated transgenic mice, pooled from two independent cohorts. Data shown as the mean ± SEM. Significance was determined by an unpaired t-test with Welch's correction. C) Representative T2-weighted image by MRI of tumor-bearing *Pten*$^{pce-/-}$/*Trp53*$^{pce-/-}$/*TMPRSS2-ERG*$^{pce+}$ mice treated with vehicle or 0.6 mg Lupron subcutaneously every 28 days for two cycles, starting 4 months after tamoxifen administration. D) Percent of mice progressing on Lupron, as measured by doubling tumor volume post treatment (>2-fold) or below this threshold (<2-fold). n=3-5 mice per group, with significance determined by Fisher's exact test. E) PTE-24 cells were treated with 1 µM GDC-941 for 72 hrs and phase contrast images were acquired.

**Supplemental Figure 2, Related to Figure 2.** A) Immune repertoire of orthotopic PTE-82 tumors following treatment with Lupron depot or αCSF-1. 17 days post implant, tumors were excised and single cell suspensions were processed for flow cytometry. B) Gene set enrichment analysis (GSEA) following whole tumor RNA sequencing. The normalized enrichment score is shown comparing tumors from mice treated with Lupron versus an IgG control. Data reflects 3 tumors per group from a single experiment. C) Volcano plots of gene expression changes between αCSF-1 or Lupron and the IgG control. The decreases in *Tmprss2, Abca1,* and *Cd36* expression are highlighted. Significant changes shared between the αCSF-1 and Lupron groups are shown in a Venn diagram to the right. D) Example extracted ion chromatograms for 9 stable isotope-labeled standards for androgens. Each ion chromatogram is labeled with the abbreviation for the androgen and the description of the stable isotopes incorporated into the standards (i.e. Carbon-13 or Deuterium). These data from parallel reaction monitoring of each precursor ion indicate the sum of ion signal intensity for the fragment ions listed in the table (lower right).

**Supplemental Figure 3, Related to Figure 3.** A) Correlation between percent F4/80 positive cells and AR nuclear staining in orthotopic PTE-82 tumors. Data represent matched averaged ROI (3-5/slide) in two serial section stained with F4/80 and AR from one cohort of mice, n=6. B) Correlation between percent F4/80 positive cells and AR nuclear staining in *Pten*$^{pce-/-}$/*Trp53*$^{pce-/-}$/*TMPRSS2-ERG*$^{pce+}$ and *Pten*$^{pce-/-}$/*Trp53*$^{pce-/-}$/*TMPRSS2-ERG*$^{pce-}$ mice at 6- and 12-month post tamoxifen. n=13-16, data pooled from two independent cohorts. C) Kernel density estimation of the AR nuclear to cytoplasmic ratio on a cell-by-cell basis following incubation of PT-09 or PTE-82 cancer cell lines alone, or in co-culture with BMDMs under androgen-deprived conditions (i.e., charcoal-stripped serum, CSS) for 72 hrs. 6-8 randomly selected images from two wells of the chamber slides were pooled for analysis. Data reflects one of at least three independent experiments. Significance determined by Mann-Whitney. D) Impact of BMDMs on the proliferation of PTE-82 cells in the presence of serial concentrations of enzalutamide. Cell proliferation was monitored using live imaging with phase contrast images acquired every 6 hrs. Data shown as the mean ± SEM from one of two independent experiments. Significance determined by two-way ANOVA.

**Supplemental Figure 4, Related to Figure 4. A)** Intergene correlation analysis of genes associated with macrophage infiltration (*CD86, CSF1R, MRC1, CD163*) and the indicated metabolic genes using **w**hole transcriptome data from the Decipher GRID$^{TM}$ registry data. B) Histogram of BODIPY-493/503 fluorescence in leukocyte populations within PTE-82 tumors. n=5, data reflects one of at least three independent experiments. C) Mean BODIPY-493/503 fluorescence intensity in myeloid populations within tumors from *Pten*$^{pce-/-}$/*Trp53*$^{pce-/-}$/*TMPRSS2-ERG*$^{pce+}$ mice, 6 months post tamoxifen. Fluorescence minus one (FMO) is shown for tumor samples. n=4, data shown as the mean ± SEM. Similar results were observed in orthotopic models and 12 months post tamoxifen in *Pten*$^{pce-/-}$/*Trp53*$^{pce-/-}$ and *Pten*$^{pce-/-}$/*Trp53*$^{pce-/-}$/*TMPRSS2-ERG*$^{pce+}$ models.

**Supplemental Figure 5, Related to Figure 5.** A) Expression of *Cyp11a1* and *Cyp17a1* in prostate cancer cell lines and BMDMs, as determined by real time PCR. n=3-5, data shown as the mean ± SEM from one experiment. B) Nuclear to cytoplasmic AR ratio of GFP$^{+}$ PTE-82 cells cultured in CSS or co-cultured with BMDMs. 1 μM BLT-1 (SCARB1 inhibitor) and 5 μg/ml αCD36 were added to co-cultures as indicated. 6-8 randomly selected images from two wells of the chamber slide were pooled for analysis. Data reflects one of two independent experiments. Significance determined by Mann-Whitney. C) PTE-82 cells were treated with the LXR modulators RGX-104 (5 μM) and SR9243 (5 μM) for 48 hrs and effects on gene expression related to cholesterol influx/efflux (*Abca1*, *Ldlr, Vldlr, Scarb1*), cholesterol *de novo* synthesis (*Acaca, Hmgcr, Sqle*), and the LXR family (*Nr1h2, Nr1H3*) were determined by real time PCR. Expression was normalized across groups and is shown as a heat map. Data represents the mean of 3 technical replicates, with significance determined by two-way ANOVA. One of two representative experiments is shown. D) PTE-82 cells were treated with 5 μM RGX-104 and 5 μM SR9243 for 48 h and uptake of pHrodo Red labeled LDL (1 μg/ml) was measured by flow cytometry after a 3 hr incubation. Data shown as the mean ± SEM of 3-5 technical replicates, with significance determined by one-way ANOVA. One of three representative experiments is shown. E) PTE-82 cells were treated with 5 μM RGX-104 and 5 μM SR9243 for 48 hrs and expression of the androgen responsive gene, Fkbp5 was determine by real time PCR. n=3, data shown as the mean ± SEM with one of two representative experiments shown.

**Supplemental Figure 6, Related to Figure 6.** A) PTE-82 tumor cells were implanted orthotopically into prostates of C57 mice, which were then treated with Lupron depot and/or αCSF-1 as indicated. Tumor volumes were measured by MRI on the indicated days. n=13-15 mice per group, data are shown as mean ± SEM, pooled from two independent experiments. Significance determined by two-way ANOVA. B) Representative T2-weighted MRI images of the mice from A.

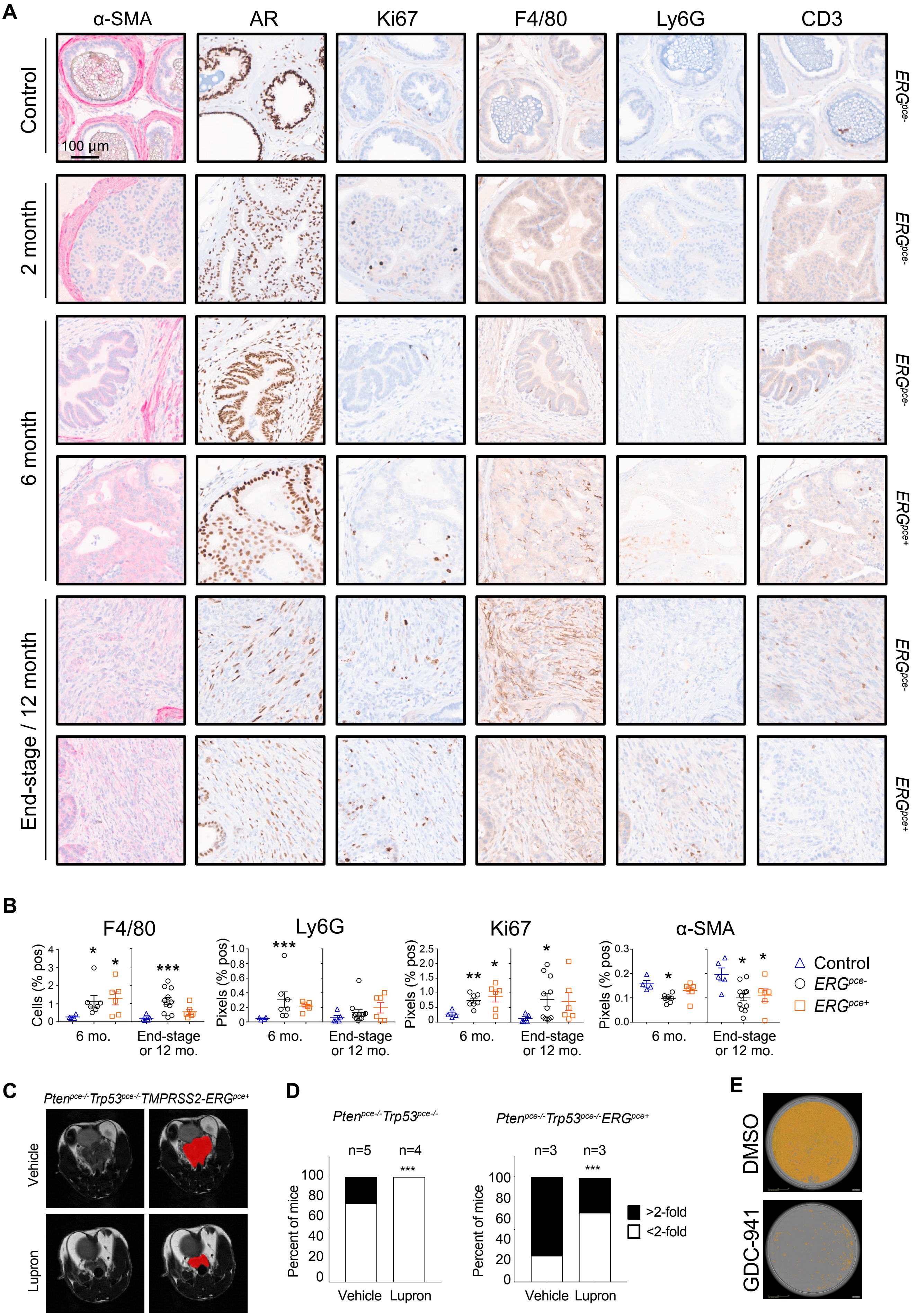
A
α-SMA
AR
Ki67
F4/80
Ly6G
CD3
Control
2 month
6 month
End-stage / 12 month
100 µm
ERGpce-
ERGpce+
B
F4/80
Ly6G
Ki67
α-SMA
Cells (% pos)
Pixels (% pos)
6 mo.
End-stage or 12 mo.
Control
ERGpce-
ERGpce+
C
Ptenpce-/- Trp53pce-/- TMPRSS2-ERGpce+
Vehicle
Lupron
D
Ptenpce-/- Trp53pce-/-
Ptenpce-/- Trp53pce-/- ERGpce+
n=5
n=4
n=3
n=3
Percent of mice
Vehicle
Lupron
>2-fold
<2-fold
E
DMSO
GDC-941

**Figure S1**

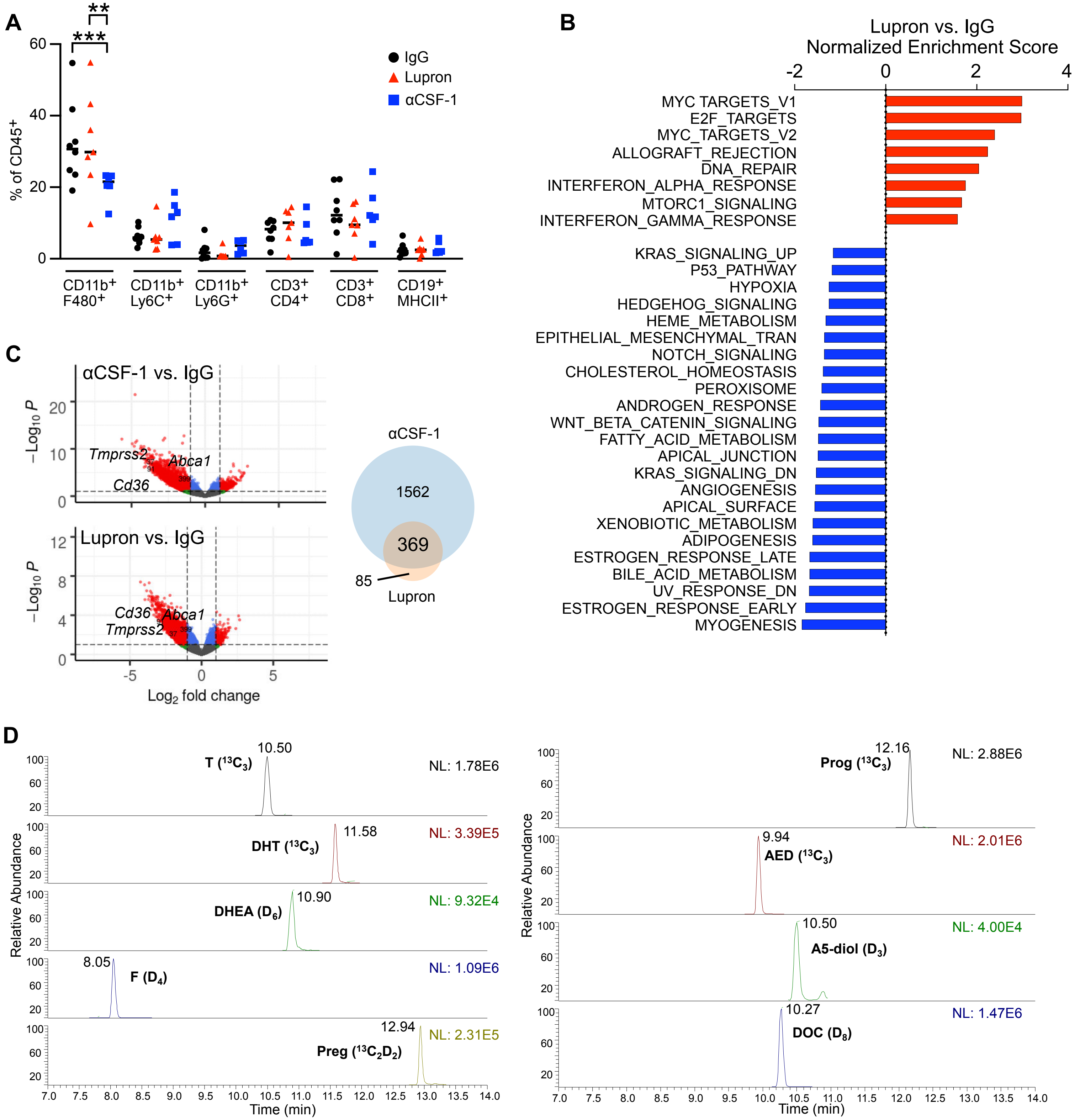


| Compound | Precursor Ion *m/z* (Ion Type) | Fragment Ion *m/z* Values |
|---|---|---|
| T ($^{13}C_3$) | 292.2263 $[M+H]^+$ | 100.0749, 112.0749 |
| DHT ($^{13}C_3$) | 294.2419 $[M+H]^+$ | 159.1168, 218.1895, 258.2208 |
| DHEA ($D_6$) | 277.2433 $[M-H_2O+H]^+$ | 219.2014, 259.2327 |
| F ($D_4$) | 367.2417 $[M+H]^+$ | 97.0649, 121.0648 |
| Preg ($^{13}C_2D_2$) | 303.2562 $[M-H_2O+H]^+$ | 161.1325, 285.2456 |
| Prog ($^{13}C_3$) | 318.2419 $[M+H]^+$ | 100.0749, 112.0749 |
| AED ($^{13}C_3$) | 290.2106 $[M+H]^+$ | 100.0749, 112.0749 |
| A5-diol ($D_3$) | 276.2401 $[M-H_2O+H]^+$ | 159.1169, 258.2208 |
| DOC ($D_8$) | 339.2770 $[M+H]^+$ | 100.0836, 113.0899 |

**Figure S2**

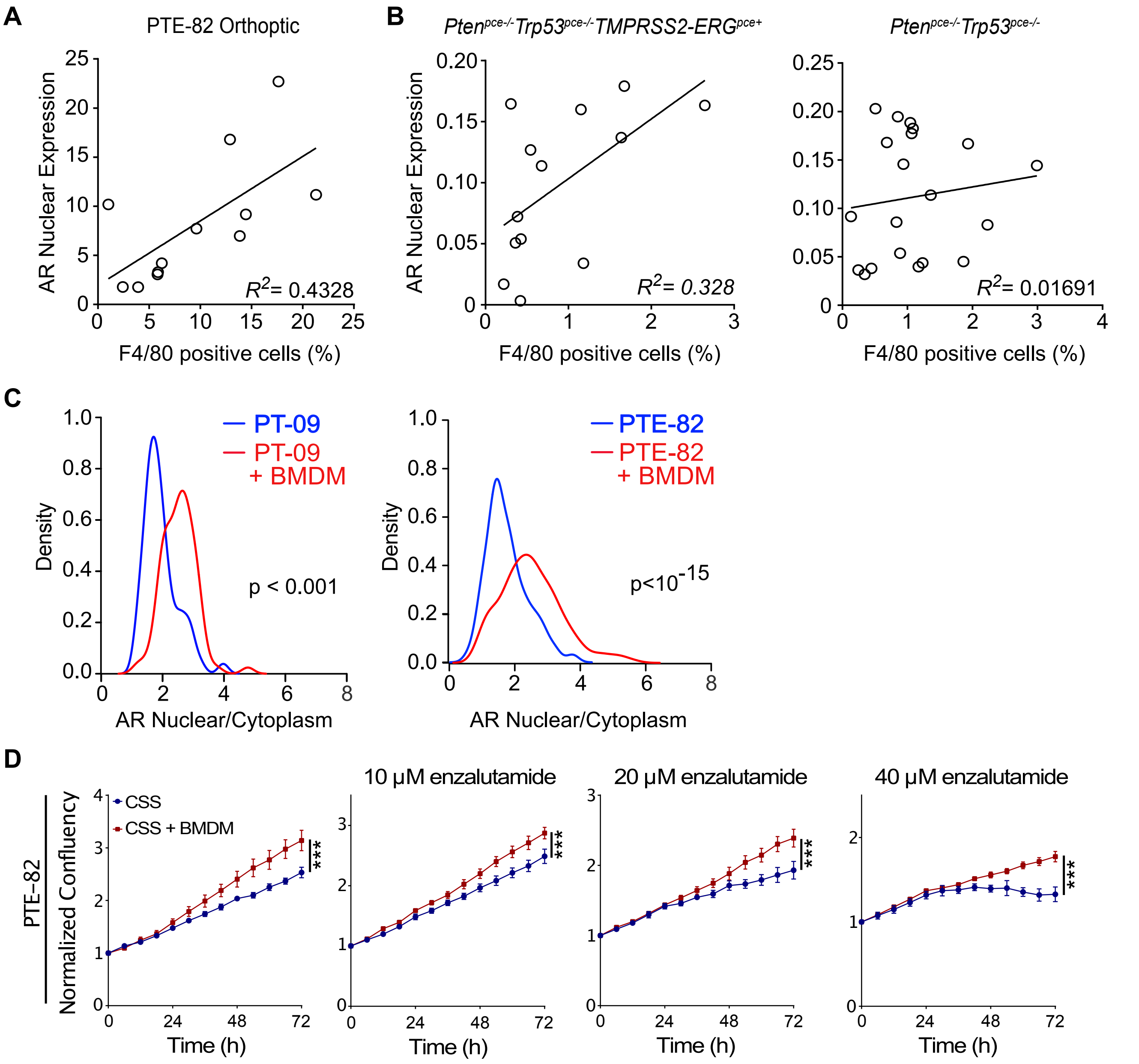
A
PTE-82 Orthoptic
AR Nuclear Expression
F4/80 positive cells (%)
R2= 0.4328
B
Ptenpce-/-Trp53pce-/-TMPRSS2-ERGpce+
R2= 0.328
Ptenpce-/-Trp53pce-/-
R2= 0.01691
C
PT-09
PT-09 + BMDM
Density
p < 0.001
AR Nuclear/Cytoplasm
PTE-82
PTE-82 + BMDM
p<10-15
D
PTE-82
Normalized Confluency
CSS
CSS + BMDM
10 μM enzalutamide
20 μM enzalutamide
40 μM enzalutamide
Time (h)
***

Figure S3

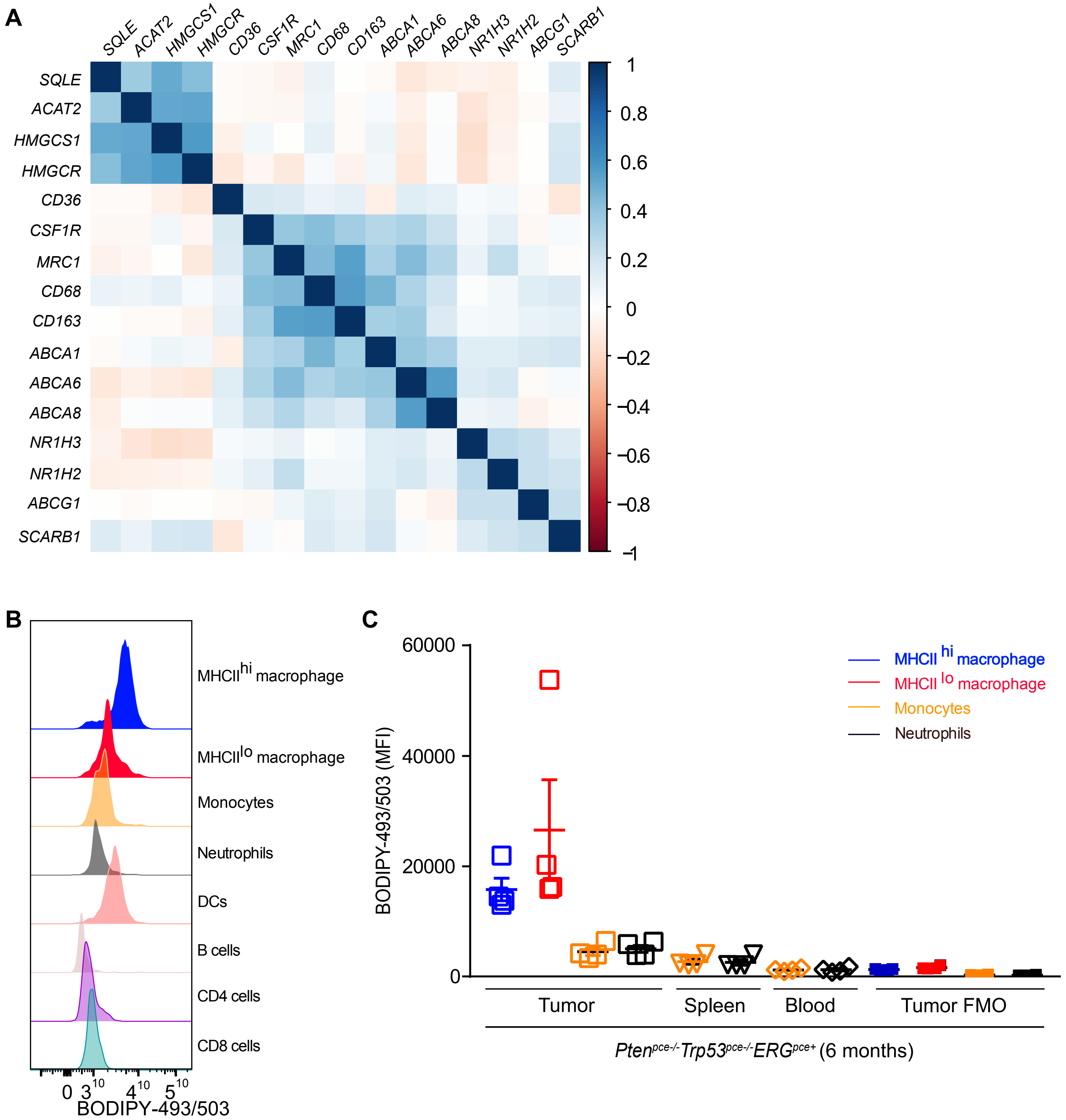

A
SQLE
ACAT2
HMGCS1
HMGCR
CD36
CSF1R
MRC1
CD68
CD163
ABCA1
ABCA6
ABCA8
NR1H3
NR1H2
ABCG1
SCARB1
1
0.8
0.6
0.4
0.2
0
−0.2
−0.4
−0.6
−0.8
−1
B
MHCII hi macrophage
MHCII lo macrophage
Monocytes
Neutrophils
DCs
B cells
CD4 cells
CD8 cells
BODIPY-493/503
C
BODIPY-493/503 (MFI)
60000
40000
20000
0
MHCII hi macrophage
MHCII lo macrophage
Monocytes
Neutrophils
Tumor
Spleen
Blood
Tumor FMO
Pten pce-/- Trp53 pce-/- ERG pce+ (6 months)


Figure S4

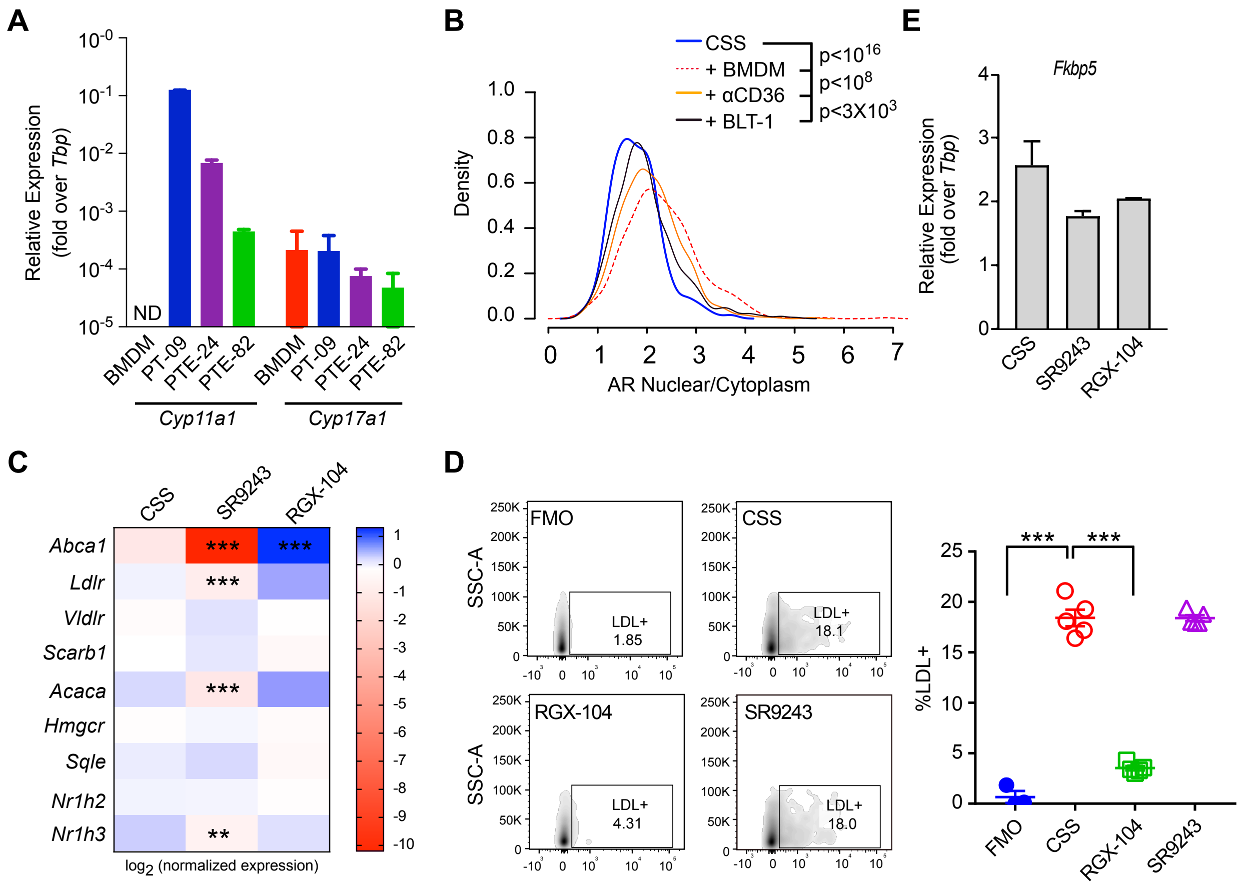
A
Relative Expression (fold over Tbp)
ND
BMDM
PT-09
PTE-24
PTE-82
Cyp11a1
Cyp17a1
B
CSS
+ BMDM
+ αCD36
+ BLT-1
p<10^16
p<10^8
p<3X10^3
Density
AR Nuclear/Cytoplasm
E
Fkbp5
CSS
SR9243
RGX-104
C
Abca1
Ldlr
Vldlr
Scarb1
Acaca
Hmgcr
Sqle
Nr1h2
Nr1h3
log2 (normalized expression)
D
SSC-A
FMO
LDL+ 1.85
LDL+ 18.1
LDL+ 4.31
LDL+ 18.0
%LDL+
***


**Figure S5**

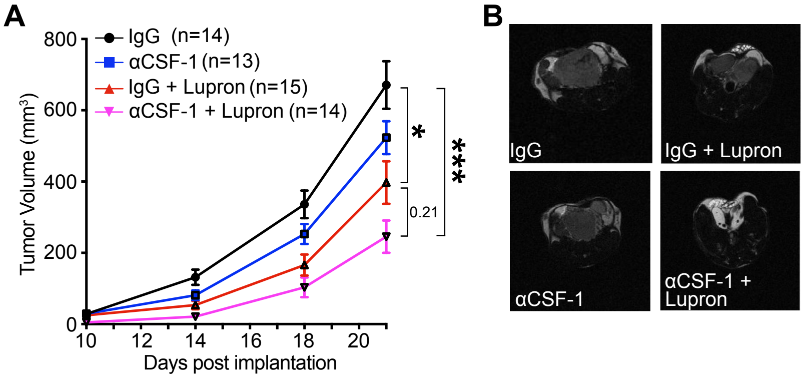
A
800
600
400
200
0
Tumor Volume (mm³)
10
12
14
16
18
20
Days post implantation
IgG (n=14)
αCSF-1 (n=13)
IgG + Lupron (n=15)
αCSF-1 + Lupron (n=14)
*
***
0.21
B
IgG
IgG + Lupron
αCSF-1
αCSF-1 + Lupron

Figure S6
